# Crowdsourcing neuroscience: inter-brain coupling during face-to-face interactions outside the laboratory

**DOI:** 10.1101/822320

**Authors:** Suzanne Dikker, Georgios Michalareas, Matthias Oostrik, Amalia Serafimaki, Hasibe Melda Kahraman, Marijn E. Struiksma, David Poeppel

## Abstract

When we feel connected or engaged during social behavior, are our brains in fact “in sync” in a formal, quantifiable sense? Most studies addressing this question use highly controlled tasks with homogenous subject pools. In an effort to take a more naturalistic approach, we collaborated with art institutions to crowd-source neuroscience data: Over the course of 5 years, we collected electroencephalogram (EEG) data from thousands of museum and festival visitors who volunteered to engage in a 10-minute face-to-face interaction. Pairs of participants with various levels of familiarity sat inside the Mutual Wave Machine—an art/science neurofeedback installation that uses Brain-Computer Interface technology (BCI) to translate real-time correlations of each pair’s EEG activity into light patterns. Because such inter-participant EEG correlations are prone to noise contamination, in subsequent offline analyses we computed inter-brain synchrony using Imaginary Coherence and Projected Power Correlations, two synchrony metrics that are largely immune to instantaneous, noise-driven correlations. When applying these methods to two subsets of recorded data with the most consistent protocols, we found that pairs’ trait empathy, social closeness, engagement, and social behavior (joint action and eye contact) consistently predicted the extent to which their brain activity became synchronized, most prominently in low alpha (∼7-10 Hz) and beta (∼20-22 Hz) oscillations. These findings support an account where shared engagement and joint action drive coupled neural activity and behavior during dynamic, naturalistic social interactions. To our knowledge, this work constitutes a first demonstration that an interdisciplinary, real-world, crowdsourcing neuroscience approach may provide a promising method to collect large, rich datasets pertaining to real-life face-to-face interactions. Additionally, it is a demonstration of how the general public can participate and engage in the scientific process outside of the laboratory. Institutions such as museums, galleries, or any other organization where the public actively engages out of self-motivation, can help facilitate this type of citizen science research, and support the collection of large datasets under scientifically controlled experimental conditions. To further enhance the public interest for the out-of-the-lab experimental approach, the data and results of this study are disseminated through a website tailored to the general public (wp.nyu.edu/mutualwavemachine).

## INTRODUCTION

Laboratory research is widely assumed to provide foundational insights into how our brains process information on an everyday basis. However, this model has not been systematically tested: we rarely, if ever, conduct our research in real-world, everyday contexts (Matusz et al. 2019b; Shamay-Tsoory and Mendelsohn 2019). At the same time, an increasing number of studies emphasize the importance of face-to-face social interaction to our physical and mental wellbeing (e.g., (Kross et al. 2013). For example, eye contact has long been recognized as a vital aspect of healthy cognition and cognitive development (e.g., (Tomasello and Others 1995), by highlighting cues that allow people to coordinate social behavior (Sebanz, Bekkering, and Knoblich 2006). To arrive at a more comprehensive understanding of the brain basis of social interaction, then, measuring communication ‘live’ is vital: realistic human interactions are more complex and more richly coupled across participants/brains than can be captured in canonical laboratory experiments.

Here, we aimed to identify neural correlates of real-world face-to-face social interactions in a large population of participants recruited outside of the traditional research subject pool (typically university undergraduate students). The homogeneity of scientific study participants is increasingly considered problematic with respect to the generalizability of research findings (Henrich, Heine, and Norenzayan 2010; Falk et al. 2013; LeWinn et al. 2017). One option is to make more of an effort to bring participants from the general public to the laboratory; another possible solution is to bring the laboratory to the public. In this work, we provide a methodological proof of concept for the second model: we show that it is feasible to conduct large-scale neuroscience research ‘in the wild’ while maintaining rigor in terms of both analysis and interpretation.

We capitalized on recent real-world social neuroscience research (Matusz et al. 2019a; Dikker, Montgomery, and Tunca 2019; Dikker et al. 2017; Bevilacqua et al. 2019; Bhattacharya 2017; Parada and Rossi 2017), mobile electroencephalography (EEG) technology (Debener et al. 2012; Gwin, Gramann, and Makeig 2010), brain-computer-interfaces (Brunner et al. 2015; Minguillon, Lopez-Gordo, and Pelayo 2017), and the so-called “interactive turn” in social neuroscience (De Jaegher, Di Paolo, and Gallagher 2010). In recent years, there has been a surge of studies that compare brain activity *between* participants instead of using a stimulus-brain approach (e.g., (Dumas et al. 2010; Hasson et al. 2004); for reviews see e.g., (Fabio Babiloni and Astolfi 2014; Hasson et al. 2012; Liu et al. 2018; Sänger, Lindenberger, and Müller 2011). Concretely, we used a ‘crowdsourcing neuroscience’ approach in which, over the course of five years, museum and festival visitors were invited to participate in research as part of their audience experience. We identified a set of characteristics that were deemed socially relevant (social closeness, social behavior, mental states, and personality traits) and asked whether these attributes affected the similarity of brain activity between two people during naturalistic face-to-face interaction (often referred to as brain-to-brain synchrony or inter-brain synchrony; operationalized below). Crucially, we not only sought to identify such factors, but also whether they can be dissociated at the neural level, specifically with respect to different characteristics of brain oscillations. Our experimental question was made tangible and enticing to the audience as follows: “When are your brainwaves literally “on the same wavelength”?”

Recent research has identified a number of predictors of inter-brain synchrony (e.g., (Hasson et al. 2004; Nummenmaa et al. 2012; Dikker et al. 2017; Bevilacqua et al. 2018; Pérez et al. 2018; Parkinson, Kleinbaum, and Wheatley 2018; Dikker et al. 2014; Stephens, Silbert, and Hasson 2010; Konvalinka et al. 2014; Astolfi et al. 2010). Figure 1 illustrates the factors under investigation here.

**Figure 1.**
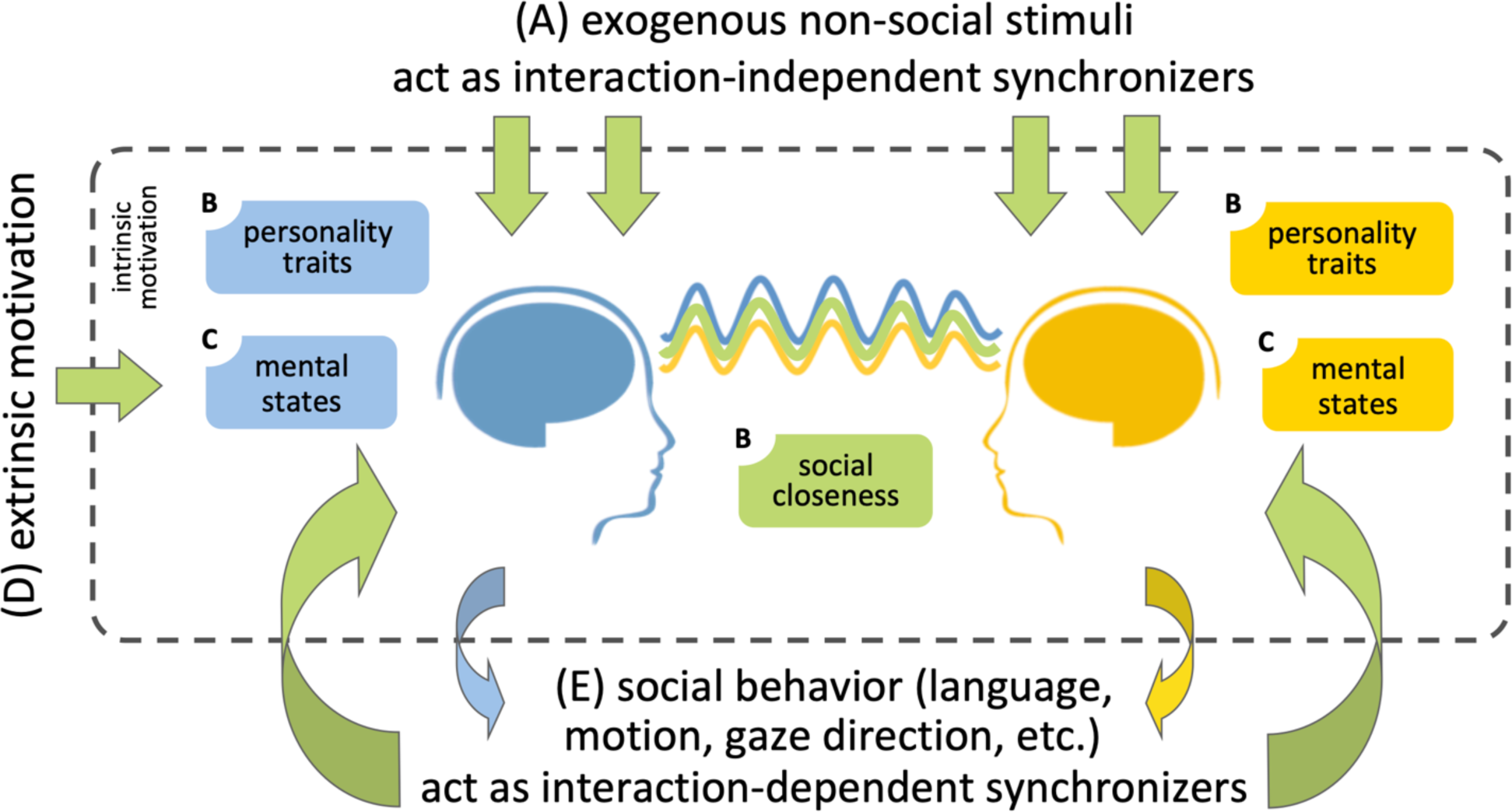
Different possible sources of inter-brain synchrony. **(A)** External non-social stimuli (top) and **(E)** social behavior (bottom) provide exogenous sources of shared stimulus entrainment and interpersonal social coordination, respectively, leading to similar brain responses, i.e., inter-brain synchrony. **(B)** Both individuals’ social closeness and personality traits (e.g., affective empathy) are expected to affect their social engagement during the interaction, and thus the extent to which their brain responses become synchronized. **(C)** participants’ mental states (e.g., focus) are similarly expected to affect participants’ engagement with each other, intrinsically (endogenously) motivate participants to make an effort to connect to each other. **(D)** Such engagement can be “boosted” via extrinsic motivation, which should subsequently lead to increased inter-brain synchrony.

It is widely established that brain activity becomes synchronized between people when they listen to or watch the same stimulus merely due to its physical characteristics (contrast, color, volume variations; Figure 1a). Another (partially) exogenous driver of inter-brain coupling is synchronized movement through joint action (e.g. (Dumas et al. 2010)). Socially-induced behavioral synchrony (Figure 1e) is prevalent throughout our everyday interactions: consider pedestrians navigating sidewalk traffic, conversations, a tango dance, a musical duet. Face-to-face interactions require tight spatio-temporal coordination between their participants at cognitive (Pickering and Garrod 2013), perceptual (Kang and Wheatley 2017), and motoric levels (D. C. Richardson, Dale, and Tomlinson 2009). Such interpersonal rhythmic coordination occurs spontaneously (M. J. Richardson et al. 2007) and is subject to individual differences: people with a prosocial orientation tend to synchronize more (Lumsden et al. 2012), and children with Autism Spectrum Disorder do not engage in spontaneous rhythmic movement synchronization with others (Marsh et al. 2013). Perhaps most importantly, synchronized joint action is predictive of how the interaction is experienced. For example, therapists and patients who exhibit more synchronized motion during a therapy session, report higher therapeutic satisfaction (Ramseyer and Tschacher 2011; Koole and Tschacher 2016; Koole et al., n.d.). Synchronous biological rhythms have also been linked to social behavior in a meaningful way, ranging from heart rate and respiration (Noy, Levit-Binun, and Golland 2015; Müller and Lindenberger 2011; Thorson, West, and Mendes 2018; Waters et al. 2017), to brain responses: For example, synchronous resting state fMRI activity between children and their caregivers is predictive of their relationship (Lee, Miernicki, and Telzer 2017), and friends show more similar neural responses to video clips (Parkinson, Kleinbaum, and Wheatley 2018).

While evoked neural activity is observed across cortex, it is typically most strictly time-locked to the stimulus--and thus also between participants--in sensory cortex (David, Kilner, and Friston 2006). Inter-brain coupling arising from internal mental models, in contrast, is arguably rooted in higher brain areas and involves top-down processes which fuse the exogenous stimulus information with such endogenous models of the world. We pose that socially relevant factors (Figure 1b-e) operate on such brain endogenous models involving high-level inferences. We thus focused our analysis on non-instantaneous co-variations in pairs’ brains: while we recognize that coupling in higher brain areas can also be instantaneous, we reasoned that non-instantaneous inter-brain coupling is less likely to arise from purely stimulus-related factors and thus more likely to stem from socially-relevant factors.

Such socially relevant factors include joint action (Figure 1e) as discussed above, but also social personality traits (e.g., affective empathy) and pairs’ social closeness, which we predicted would both affect pairs’ mutual social engagement, and thus the extent to which their brain responses become synchronized. Past research has already demonstrated that interpersonal factors affect inter-brain coupling. For example, collaborative attitudes lead to higher inter-brain coupling than competitive behavior (Cui, Bryant, and Reiss 2012) as do social closeness and empathic personality (e.g., (Kinreich et al. 2017; Goldstein et al. 2018; Dikker et al. 2017; Bevilacqua et al. 2019).

Mental states such as engagement/focus level (Figure 1c) have also been shown to affect inter-brain coupling (Dumas et al. 2012; Dikker et al. 2017; Goldstein et al. 2018; Dalton et al. 2005; Scott-Van Zeeland et al. 2010; Kylliäinen et al. 2012; Petroni et al. 2017; Bevilacqua et al. 2019). Such engagement can arguably be “boosted” via extrinsic motivation (Figure 1d), which should subsequently lead to increased inter-brain coupling.

To test how socio-behavioral factors during face-to-face interaction relate to inter-brain coupling, we developed the Mutual Wave Machine (Dikker, Montgomery, and Tunca 2019), a dome-like BCI/neurofeedback environment that immerses pairs of participants in a real-time audio-visual (AV) reflection of the extent to which their EEG signals are instantaneously correlated in one or more frequency bands (Figure 2).

**Figure 2.**
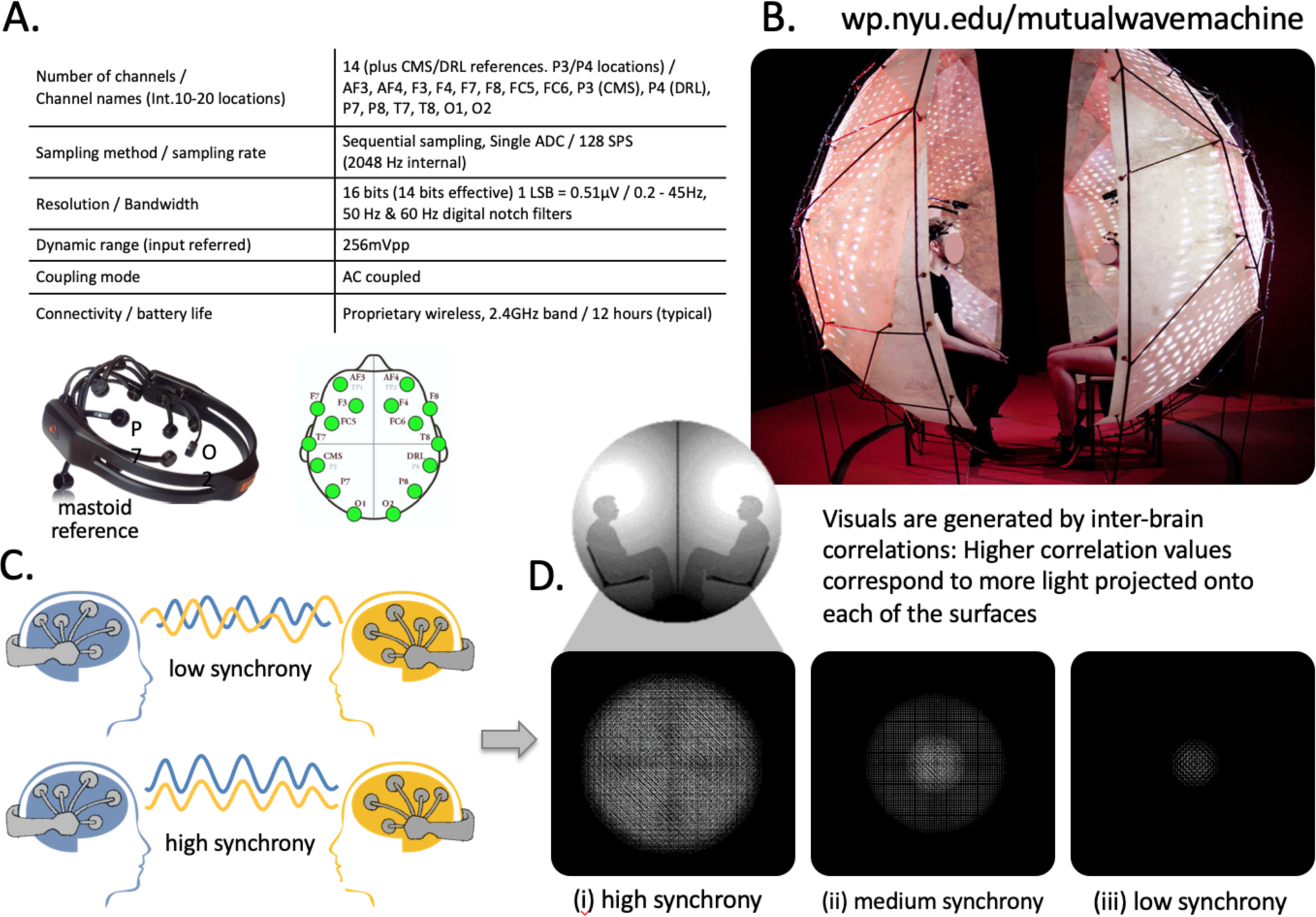
The Mutual Wave Machine. **(A)** Hardware specifications of the EMOTIV EPOC EEG headset, an image of the EMOTIV EPOC headset (side view), and a top-view of the electrode locations (note that electrode placement may vary considerably between participants, see e.g., Dikker et al., 2017); **(B)** A pair of participants inside the Mutual Wave Machine experiencing real-time inter-brain coupling A/V feedback. **(C/D)** Inter-brain correlations between two participants wearing wireless EEG headsets were computed in real time. Higher inter-brain correlation values correspond to more light projected on each of the surfaces, with the focus point behind each participant’s head. (See text for details; for the offline inter-brain coupling computations see Figure 3).

We hypothesized that such an environment would motivate participants to remain socially engaged with each other for the duration of the interaction (Figure 1d). To test this, we asked whether explicitly informing participants that the audio-visual (A/V) environment reflected their ongoing correlated EEG signal resulted in a “self-fulfilling prophecy” of sorts: If you think you are receiving real-time audio-visual feedback about how in sync you are with your partner, will that actually positively affect your ongoing inter-brain coupling?

As detailed in the Methods, it is important to note that ‘brain synchrony’ as reflected in the Mutual Wave Machine is likely very prone to noise contamination, rendering it difficult to draw meaningful conclusions about participants’ veridical brain synchrony based on their real-time A/V environment. Participants were explicitly informed of this limitation and were told that a subsequent offline scientific analysis would be applied to their data. A second limitation is that the real-time brain synchrony involved the computation of instantaneous correlations between narrow-band versions of short portions of the EEG recorded signals. As a result, high correlation values due to chance are more likely than when broadband signal correlations are used (see Methods for a more detailed description). These limitations, together with the fact that the correlational approach that was implemented here is not commonly used in hyperscanning research, decreases the likelihood that the neurofeedback is socially significant.

To circumvent these issues, we used two different metrics to compute non-instantaneous inter-brain coupling: *projected power correlations* and *imaginary coherence*. While imaginary coherence provides a quantitative characterization of whether oscillations within specific frequency ranges show consistent phase relationships between pairs of participants; projected power correlations quantify the extent to which power in a specific frequency co-fluctuates between pairs. Second, as explained in detail in the Methods, projected power correlations and imaginary coherence are especially suited for noisy recording environments, such as museums (Cruz-Garza et al. 2017), because the common signal between individual recordings (or: instantaneous co-fluctuations) is removed before computing inter-brain coupling. In other words, any electrical noise or sensory cortex activity driven by strong audio-visual input can only minimally affect the estimated inter-brain coupling. This is especially important in the current setup because participants were surrounded by electronic equipment and strong audio-visual stimulation originating from the art exhibition. We additionally circumvented spurious effects of environmental noise on inter-brain coupling by adopting a strict brain-behavior correlational analysis approach.

To summarize, in this study pairs of participants interacted semi-naturally while seated facing each other, allowing us to investigate how the extent to which brain activity becomes synchronized between dyads during face-to-face social interaction relates to participants’ (a) relationship (relationship duration and social closeness; (Aron, Aron, and Smollan 1992), (b) affective personality traits (Personal Distress and Perspective Taking; (Davis and Others 1980), (c) mental states (positive affect, negative affect, and focus; (Watson and Clark 1994), and (d) motivation (real-team audio-visual brain synchrony feedback; (Dikker, Montgomery, and Tunca 2019). To address these questions, we used a BCI setup that allowed us to collect neural activity using portable electroencephalography (EEG) from large numbers of museum and festival visitors at a wide range of recording sites (see Table S1 for a list of recording sites and the number of participants for each site). This interactive art/science neurofeedback ‘laboratory’ enabled us to explore the limits and opportunities afforded by conducting human social neuroscience research outside of the traditional laboratory context.

## METHODS

### Data Recording and Real-time Analysis

Pairs of participants sat inside the Mutual Wave Machine (Figure 2b) while we recorded their brain activity using 14-channel portable EMOTIV EPOC wireless EEG headsets (see Figure 2a for specifications, and e.g., (Debener et al. 2012; Dikker et al. 2017), for a validation). ‘Brainwave synchrony’ (in this case, instantaneous band-limited correlations, described below) was translated into light patterns that were projected onto the surface of the installation (Figure 2b-d, see wp.nyu.edu/mutualwavemachine for footage of the installation/setup) using custom-software developed in the C++ based OpenFrameworks library (openframeworks.cc).

The exact data processing pipeline that was used to compute the inter-EEG correlations that fed into the visualizations varied between the locations listed in Table S1. For the BENAKI and LOWLANDS datasets described here, the following protocol was used. A moving-window of 6 seconds of data (6 * 128 samples) was selected for real-time analysis approximately 30 times per second (the exact window-step size varied slightly as a function of the buffer size of incoming samples, which typically ranged between 4-8 samples per buffer). EEG data streams from the two participants’ headsets were synchronized using a 5-second window, based on the time that samples were received by the analysis computer, and then filtered into typical frequency bands using FFTW (www.fftw.org; delta: 1-4 Hz; theta: 4-7 Hz; alpha: 7-12 Hz; beta: 12 -30 Hz). Within each frequency range, a sub-window was used to calculate Pearson correlation coefficients for each pair of sensors between headsets (n x n sensors; delta: 3 * 128 window, 0.7 * 128 offset; theta: 2 * 128, 0.6 * 128 offset; alpha: 1.1 * 128 window, 0.5 * 128 offset; beta: 0.6 * 128 window, 0.4 * 128 offset). Both the average across all r-values for all sensor pairs and the highest r-value among all sensor pairs contributed to the correlation value for each frequency band (50/50). The four scores were then fed into a visualization algorithm in which four separate moiré patterns were created, growing in a circular motion from the center of each sphere as a function of the synchrony value in each frequency band, ranging between 0-1: a score of 1 was screen filling (Figure 2d-ii) and no visuals were projected if the value was 0 (Figure 2d-iii).

Although this setup makes “brain synchrony” intuitive to the general public, it is highly unlikely that these instantaneous band-limited correlations map onto inter-brain coupling between the participants in a meaningful way. The main reason is that in a noisy environment, such as a museum, instantaneous band-limited correlations between the 2 EEG devices are likely to be dominated by shared noise rather than shared social events. Additionally, when correlating 2 short narrow-band signals, correlation values can be artificially inflated because of pure chance. This inter-brain coupling analysis was used for ease of computation and to our knowledge has not been employed to assess meaningful EEG inter-brain analysis.

As described in the Procedure, participants were made aware of the caveats and limitations of the visualization and were discouraged to draw any meaningful conclusions based on their experience. Instead, they were told that for analysis, their brain coupling would be recomputed later, offline, after removing motion artifacts and bad channels. They were told that once the analysis is done, findings would be made available on a public website.

### Participants & locations

EEG and questionnaire data were collected from a total of 4,800 people across 14 different sites (see Table S1). For the purposes of this paper, data were analyzed only from two sites. The Benaki Art Museum (BENAKI; Athens, Greece), where the Mutual Wave Machine was set up as part of the Marina Abramovic Institute exhibition AS ONE (mai.art/as-one-content/2016/2/29/presenting-as-one), provided our most comprehensive and consistent dataset (1,568 participants) and provided the best recording conditions: the Mutual Wave Machine was exhibited for 2 months in the same location, the data were collected by a well-trained and dedicated group of facilitators (see Acknowledgments), and there was minimal environmental distraction (at some of the other sites, the installation was placed in a crowded space with a lot of environmental noise and distractions for both participants and experimenters). The second recording site included here is Lowlands Science (LOWLANDS), where 230 participants took part in the Mutual Wave Machine.

### Experimental procedure

Participants signed up for timeslots in advance, either individually or in pairs. EEG headsets were applied while participants completed a consent form (following Utrecht University Institute of Linguistics study protocol) and pre-experiment questionnaire (see below for details). After setup, participants were led to the Mutual Wave Machine, where they received further instructions.

In the BENAKI dataset, a subset of participants (n = 534) was explicitly told that the light patterns reflected brain-to-brain synchrony (explicit synchrony A/V), while another group (n = 498) was not (non-explicit synchrony A/V). In the LOWLANDS dataset, all pairs were told the purpose of the work was to investigate the relationship between inter-brain synchrony and “feeling in sync”, but half of the participants were assigned to a sham A/V condition. For the latter group, the visualizations were randomly generated instead of reflecting the true correlated EEG signal. An additional difference in the LOWLANDS dataset was that, after the experience, participants were asked to list which “strategies” they used to try to connect to each other and increase brain synchrony.

Participants were encouraged to be mindful of their movements and were told that too much movement would create motion artifacts that could distort the signal and make it impossible to detect actual brainwaves in the EEG signal. To illustrate this, participants were shown the effects of jaw clenching, laughing, and blinking in their own raw EEG trace during setup.

After the experience, participants were led back to the setup station to fill out an additional set of questions. They were told that their data would be scientifically analyzed offline and that results would be made available on a publicly accessible web page (wp.nyu.edu/mutualwavemachine). The experiment took approximately 20 minutes including setup and debriefing.

## Materials

To investigate which factors drive inter-brain coupling during spontaneous face-to-face interaction, participants were asked to fill out a series of questions designed to address their (a) relationship to each other, (b) affective personality traits, and (c) affective mental states.

a. **Relationship metrics** were administered before the experience only, and included four variables. (i) As an index of **Relationship Duration (a trait measure)**, participants were given 6 choices to indicate how long they knew each other (varying from 1 = Strangers to 6 = 10+ years). (ii) **Social Closeness (a state measure)** between pairs was measured using the Inclusion of the Other in the Self Scale (Aron, Aron, and Smollan 1992), in which participants are presented with 6 Venn diagrams where circles ‘Self’ and ‘Other’ overlap to varying degrees, and are asked which best applies to his/her relationship to the other.
b. To quantify socially relevant **affective personality traits**, participants completed a revised version of the Interpersonal Reactivity Index during setup (IRI; (Davis and Others 1980), including the subscales **Perspective Taking** (e.g., “When I’m upset at someone, I usually try to “put myself in his shoes” for a while.”), and **Personal Distress** (e.g., “When I see someone who badly needs help in an emergency, I go to pieces”). The questionnaire consisted of 14-items answered on a five-point Likert scale ranging from “Does not describe me well” to “Describes me very well”.
c. Mental state metrics were measured both before and after the experience (Pre and Post) using a shortened version of the Positive and Negative Affect Schedule (PANAS-X; (Watson and Clark 1994). The questionnaire consisted of 20 items, with 10 items measuring Positive Affect, of which three items measured **Focus** (e.g., attentive, alert). The items were rated on a five-point Likert Scale, ranging from 1 = *Very Slightly or Not at all* to 5 = *Extremely*.

### Offline Analyses

#### Preprocessing pipeline

The initial pool of data consisted of EEG recordings of 1,568 participants in the BENAKI dataset, and 230 participants in the LOWLANDS dataset (see above and Table S1).

First, raw datasets with problems due to headset or recording software malfunction were identified and discarded. For the BENAKI dataset, 28 pairs were rejected because for one or both participants, less than 5 minutes of raw data were available, likely due to false starts or other recording issues (out of 10 minutes total; mean duration of all pairs = 551.6 s). An additional 117 pairs were rejected because of intermittent data loss (93 pairs), data repetition (34 pairs), or drift between the two headsets (3 pairs), factors that would render inter-brain coupling analyses unreliable.

Next, we identified physiological artifacts or channel specific hardware artifacts. The EEG headset provides binary variables, sampled at the same rate as the data, which mark blinks or vertical/horizontal eye movements. All the instances for which these three flag variables were true were marked as artifacts, as well as 50 msec before and 250 msec after such instances. Subsequently, four different types of artifacts were identified using the Fieldtrip toolbox (Oostenveld et al. 2011). *<u>Signal</u> <u>Jumps</u>* are sudden (step-like) increases or decreases in the recorded electric field, usually attributed to the amplifier electronics. *<u>EOG-like</u> <u>Artifacts</u>* (electrooculography) were identified based on their typical band–limited characteristics (no EOG was recorded during the experiment). For the purpose of removing EOG-like artifacts, the EEG channels were band-passed by a Butterworth filter in the frequency range [1-15 Hz], then envelope time-series were derived through the Hilbert transform, and times when this envelope exceeded 4 standard deviations from its mean were marked as potential EOG artifacts. *<u>Clipping</u> <u>Artifacts</u>* are periods when data in one or more channels remain constant at a given value and is usually caused by short-term problems in the electronics. *<u>Head</u> <u>Movements</u>* were identified through a 2-axis gyroscope built into the EEG headset, providing information about the orientation of the head at each data sample. The accelerometer signal in each axis takes negative and positive values depending on the direction of movement. Significant head movements are manifested as large changes on the accelerometer signal. In each of the 2 accelerometer axes the derivative of the accelerometer signal was computed first and subsequently the mean and standard deviation of this derivative were calculated. All data points for which the derivative was 5 standard deviations above and below its mean were marked as head movement artifacts. Data instances with any of the above artifacts characteristics were marked as “bad” segments.

The EEG headset provides, at the same sampling frequency as the data, a quality flag for each of the data channels. This data quality flag is a measure of contact quality of a given electrode on the scalp during a recording, ranging from 0 to 4. Any channel with an average quality lower than 1 was identified as a noisy, “bad” channel and discarded from further analysis (still preserving some useful information).

After the bad channels were discarded from a dataset, the raw data were segmented into pseudo-trials of 1 second duration each, and all the pseudo-trials that coincided with “bad” segments, as described above, were completely removed from the data. This pseudo-trial representation was used so that in subsequent time-frequency and band-limited connectivity analysis data segments surrounded by discarded artifacts would have a duration of an integer number of seconds.

Once bad channels and bad segments were removed, one last automated preprocessing procedure was applied to further reduce the potential data contamination by artifacts. Given the nature of the electric signals emanating from the brain and measured on the scalp, the variance of the recorded data should be similar across the different channels. Channels with consistently higher variance compared to the other channels across the whole recording are most likely to contain higher noise, likely due to electrical noise and not due to environmental noise (the latter would affect all channels similarly). In order to investigate such cases, each dataset was divided into 10 equally sized segments and the variance of each channel was computed for each segment. If, across most of the 10 segments, a channel was found to consistently have a variance higher than 2 standard deviations from the average variance over all channels, it was flagged as a “bad” channel. We also examined if specific trials showed much higher variance than the average variance across all trials. As the paradigm employed here did not include any specific stimulus presentation, the variance was expected to not have big fluctuations across time but stay within relatively stable boundaries. Periods of higher variance are indicative of muscle or movement artifacts. So as a first step, all trials with a variance of larger than 3 standard deviations above the average variance across the recording in a channel were marked as artifacts. In addition to this channel-specific “bad” segment identification, a final similar analysis was performed, but this time the variance for each trial was averaged across channels. This reflected trials that have high variance consistently across all channels, probably due to higher environmental noise. Again, pseudo-trials with a variance larger than 3 standard deviations from the mean variance across the recording were marked as “bad” segments.

These “bad-channel” and “bad-segment” procedures were applied to the recordings of each of the participants in a given pair. This resulted in a cleaned dataset for each participant, comprising a set of 1-sec long pseudo-trials with gaps in the places where potential artifacts were identified. Only trials for which data were available for both participants (aligned trials) were kept for further inter-brain coupling analysis. 331 pairs had missing or incomplete questionnaire data (due to internet connectivity issues at the museum; the questionnaires were administered using an internet-based platform), leaving 307 pairs for further analysis.

The same preprocessing procedure was applied to the LOWLANDS dataset and resulted in a total of 53 pairs (out of 115) to be included for further analysis.

#### Time-frequency analysis

The raw continuous data was first demeaned and high-pass filtered at 0.5 Hz in order to remove slow fluctuations. Then all the “bad” channels and “bad” segments, defined through the methodology described above, were removed. Time-frequency analysis was performed using a Hanning-taper transformation based on multiplication in the frequency domain (Maris & Oostenveld, 2007). The investigated frequencies were between 1 and 40 Hz, in steps of 1 Hz. The time-frequency analysis was performed for all time points of the data. For each time point and given frequency, the spectral complex coefficient was computed from a data window centered at that timepoint. The length of the window was selected to be 5 periods of the frequency in question unless this exceeded 500 msec, in which case the window length was set to this maximum value. The different window lengths with regard to frequency created a frequency resolution different from the desired 1 Hz. In order to achieve this resolution, zero-padding was employed. This analysis resulted in a series of complex coefficients for each channel and frequency, with gaps where artifacts had been removed and when not enough data samples where available to fill the time-frequency window. These results were stored and used in the following connectivity analysis.

#### Inter-brain functional connectivity analysis

Inter-brain coupling can be measured in various ways. Here, it was quantified using two metrics: imaginary coherence and projected power correlations. This choice was motivated not only because of the proposed mechanisms they are argued to capture, but also because of the recording conditions. Specifically, coupling can be manifested in the fluctuations of the recorded electrical signals themselves or in the fluctuations of the power envelopes of these signals. Coherence is the most typical functional connectivity metric employed to study the former type of coupling, and power correlations are most often used for the latter. However, both metrics, especially when applied to EEG and MEG measurements, are highly contaminated by environmental noise, which is manifested as common signals with 0 phase difference across sensors and headsets. In the current study, such noise-induced zero-lag signals are even more pronounced, since the museum environment has higher noise than a typical EEG lab (e.g., via light installations on the ceiling as well as audio and video electronic equipment in the vicinity of the measurements).

To avoid such instantaneously synchronized EEG signals within pairs, which have no relevant neurocognitive interpretation, *Imaginary Coherence* (IC; coupling in EEG signal fluctuations; (Nolte et al. 2004) and *Projected Power Correlations* (PPC; coupling in the signal envelopes after the projection of one signal on the other has been removed; (Hipp et al. 2012) were employed instead (schematic in Figure 2e).

Assume there are two EEG channel recordings (*t*), *y*(*t*) whose time-frequency spectral coefficients series are *X*(*t, f*) and *Y*(*t, f*). Imaginary coherence takes two series of complex spectral coefficients and computes the cross spectral density *S*_*yy*_(*f*) between them (eq. 1.3) and the auto spectral density (power) of each of them, *S*_*xx*_(*f*), *S*_*yy*_(*f*) (eq. 1.1 and 1.2). From these, it computes the Coherency *C*_*xy*_(*f*) (eq.1.4), a complex number whose phase describes the average phase difference between the two series and whose magnitude describes how consistent this phase difference is. This complex number is decomposed in a real and an imaginary part. The real part represents how much of the magnitude is driven by instantaneous interactions and the imaginary part represents how much by lagged interactions. Imaginary Coherence *IC*(*f*) is the absolute value of the imaginary part, thus represents only lagged interactions (Nolte et al. 2004). Imaginary coherence in the frequency range investigated here (1 to 40 Hz) captures phase differences between signal fluctuations in the range of tens of milliseconds.

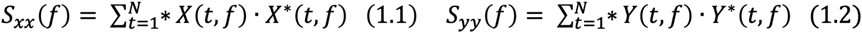

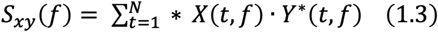

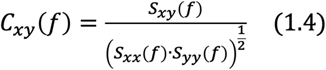

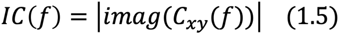

The Projected Power Correlation *PPC*(*f*), as it is called in this paper for convenience, takes two series of complex spectral coefficients and computes the correlation between their magnitudes (envelopes) after first removing the projection of one series on the other, so that all instantaneous signal fluctuations are removed before the envelope is extracted (Hipp et al. 2012). So first the projection of *Y*(*t, f*) on *X*(*t, f*) is removed, leaving only the signal orthogonal to it *Y*_┴*X*_(*t, f*)(eq. 1.6; (Hipp et al. 2012). Then the correlation between |*Y*_┴*X*_(*t, f*)| and |*X*(*t, f*)| is computed across time. The same is done also between |*X*_┴*Y*_(*t, f*)| and |*Y*(*t, f*)| (eq. 1.7). Note that *ê*_┴*Y*_ and *ê*_┴*X*_ are unit vectors representing the phases of the complex numbers and do not influence the magnitudes. Here is must be clarified that although instantaneous fluctuations of the signals are removed, the fluctuations of the envelopes of the remaining signals can still have zero phase difference, but this will not be due to a common signal present in both signals. Projected Power Correlation in the frequency range investigated here captures envelope fluctuations in the range of hundreds of milliseconds. So this analysis captures much slower processes than Imaginary Coherence.

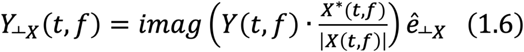

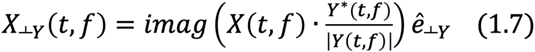

The computation of the above metrics was performed as follows. A moving window of 2-sec duration was employed to move along the series of complex spectral coefficients that were computed in the time-frequency analysis (Figure 3a). The time step this window moved at every iteration was 10 data samples (10* 1/128Hz = 70.81 msec; Figure 3b). At each iteration, all the spectral coefficients that fell within the window were selected from each matched electrode of the two headsets used in a given pair. These two time-series of spectral coefficients were then used in the computations of the connectivity metrics, and each connectivity metric was averaged across channels (Figure 3c). This was repeated for each step iteration of the moving window, so that at the end of all iterations a series of connectivity metric values across time were computed. Then the average connectivity across time was computed. This process resulted in a single value for each connectivity metric per frequency per pair (Figure 3d).

**Figure 3.**
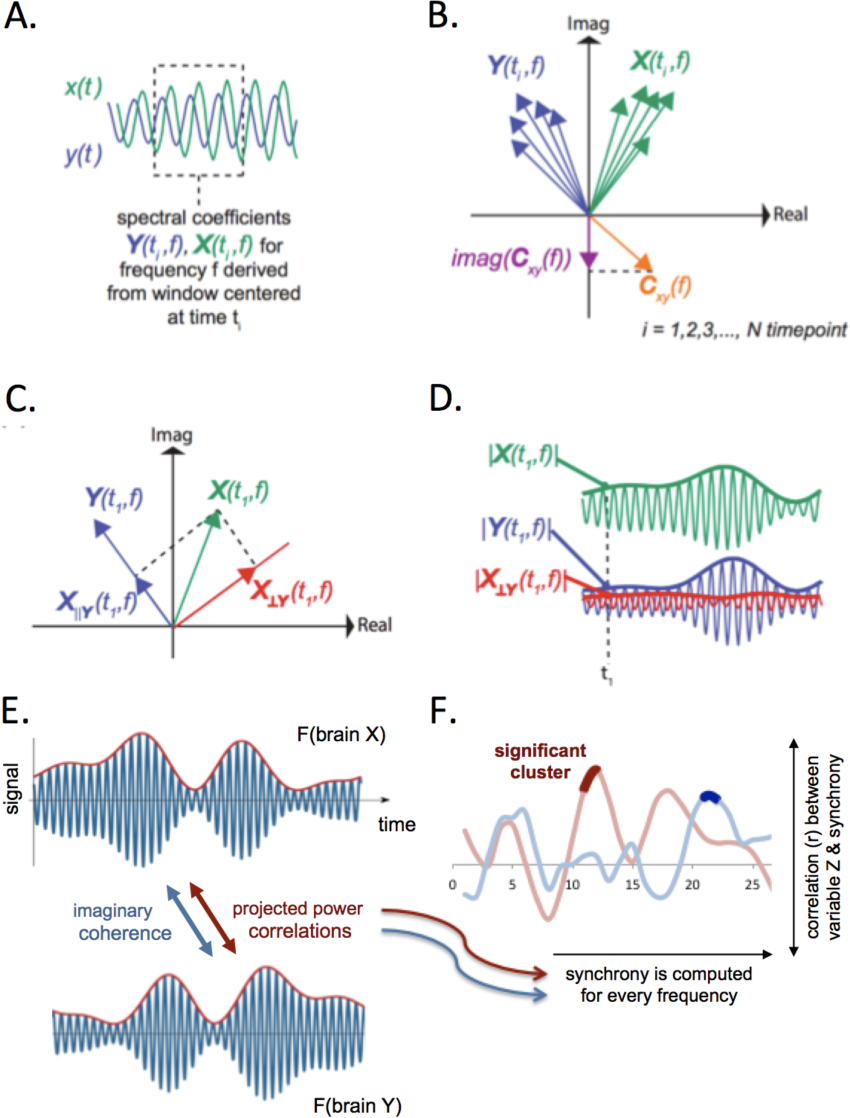
Inter-Brain functional connectivity measures. **(A)** 2-second moving window was used along each of the time-series (*x*(*t*) and *y*(*t*) correspond to the 2 different participants of a pair) of EEG recordings in order to compute the time-series of spectral coefficients *X*(*t*_*i*_, *f*), *Y*(*t*_*i*_, *f*)(one spectral coefficient per time-window *i*, per frequency *f* and participant). **(B)** These spectral coefficients for the 2 different participants can be visualized as pairs of vectors *X*(*t*_*i*_, *f*), *Y*(*t*_*i*_, *f*) at each time instance *i* (and frequency *f*). These vector pairs are then used to compute the complex coherency *C*_*xy*_(*f*), which reflects how consistent is the phase difference (angle) between the 2 participants, i.e across all spectral coefficient pairs *X, Y* (across time). Imaginary coherence constitutes only the imaginary part of coherency, reflecting only non-instantaneous phase relations (other than 0). **(C)** Projected Power Correlation: the vector of a spectral coefficient from one participant *X*(*t*_*i*_, *f*) can be decomposed into 2 orthogonal projections. One projection parallel to the vector *Y*(*t*_*i*_, *f*) of spectral coefficient of the other participant, *X*_∥*Y*_ (*t*_*i*_, *f*) and one projections perpendicular to it, *X*_⊥*Y*_ (*t*_*i*_, *f*). The parallel projection represents the part of the signal that is common between the 2 participants; as this reflects instantaneous co-fluctuations, it is removed before computing power correlations between the 2 participants. **(D)** Projected Power Correlation: The perpendicular projection *X*_⊥*Y*_(*t*_*i*_, *f*) is the part of *X*(*t*_*i*_, *f*) that is used to compute power correlations with *Y*(*t*_*i*_, *f*). **(E)** Conceptual depiction of the features that are captured by imaginary coherence and projected power correlation, computed between the EEG signals of 2 different brains X and Y. Imaginary coherence describes how consistent the phase difference is between the 2 EEG signals (EEG signals are depicted with blue). Projected power correlation describes how correlated the fluctuations are of the Envelopes of the 2 EEG signals (signal envelopes are depicted by red). **(F)** Imaginary Coherence and Projected power correlations between participants were computed at each frequency from 1 – 40 Hz and were subsequently correlated with one of the self-report variables of interest (here termed Z); a cluster analysis was used to determine significant clusters of at least 2 consecutive frequencies (dark red/blue). See text for detailed description.

The same process was also performed in the early and late halves of the data in order to investigate if engagement of the participants in the experiment increased or decreased inter-brain connectivity and if such gradients correlated with behavior.

### Correlation of inter-brain connectivity with self-report data

Once the two connectivity metrics were computed for each pair and frequency, the next step was their correlation with self-report metrics derived from the questionnaires (see Materials). This was quantified by computing Pearson correlation coefficients between a given connectivity metric and a given self-report variable for a given frequency across all pairs, accompanied by the p-value of the Pearson’s coefficient. This was repeated for all 40 frequencies for which the connectivity metric was computed. Correction for multiple comparisons was implemented as follows. With the typical significance p-value threshold of and 40 frequencies, traditional correction approaches such as Bonferroni and False Discovery rate are very conservative, highly insensitive, and fail to incorporate the fact that significant effects tend to occur in clusters along the frequency axis. Cluster-based nonparametric statistical tests based on random permutations^68^ solve the multiple comparison problem while preserving sensitivity. These non-parametric tests were used as follows. The Pearson correlation coefficient was initially computed between connectivity metric *M*(*f*) at frequency *f* and self-report variable *B*. This was repeated across all frequencies. Then correlation significance thresholds *Th*_*upper*_ and *Th*_*lower*_ were selected at the 5% and 95% percentiles of the correlation distribution for all frequencies, respectively. Then the random permutation procedure took place. The order of the self-report variable values was randomly shuffled and the correlation with the connectivity metric was repeated. All correlation values exceeding thresholds *Th*_*upper*_ and *Th*_*lower*_ were marked as significant and it was investigated if they formed clusters in adjacent frequencies. For each of these randomly significant clusters, a cluster statistic was computed. Different options were available for this average cluster statistic, such as the maximum correlation value, the average correlation value, or the size of the cluster. The extent of the cluster was chosen here because typical intrinsic oscillatory phenomena in the brain span a range of frequencies rather than single frequencies, and this should be a prominent feature in this correlational analysis too. So the randomly significant cluster with the maximum size was found and its size was stored. This random permutation procedure was repeated 500 times. At the end of this procedure, a distribution of the 500 largest random cluster sizes was formed against which all the clusters from the actual data correlation, performed at the very beginning, were compared. If an actual data cluster had a size larger than the 95% threshold of the random distribution, it was marked as significant (Figure 3f). This procedure was applied to each connectivity metric and each self-report metric.

## RESULTS

The results below quantify correlations between self-report variables and inter-brain coupling in the BENAKI dataset, unless indicated otherwise. All reported p-values survived correction for multiple comparisons using a False Discovery Rate approach (FDR; (Benjamini and Hochberg 1995)

Socially relevant trait measures, including pairs’ relationship (Aron, Aron, and Smollan 1992) and affective personality traits (Davis and Others 1980), were correlated with inter-brain coupling averaged across the entire 10-minute experience. For pairs’ focus and motivation, we instead examined changes in inter-brain coupling: the difference between the average inter-brain coupling during the second half of the experience and the first half of the experience was correlated with these state measures. Inter-brain coupling was quantified in two different ways. First, Projected Power Correlations capture band-limited power fluctuations at different oscillatory frequencies. These power fluctuations represent relatively slow processes (on the order of seconds) and represent the overall strength of neural activation. With “overall strength of neural activation” at a particular frequency, we refer to a greater number of neurons firing synchronously at a particular frequency. The larger the number of neurons firing synchronously at that frequency, the larger the magnitude of the measured electric field at this frequency and the higher the power. Note, however, that projected power correlation quantifies the level of co-fluctuation of this magnitude across time between two signals and not of the phase of firing. In addition to power fluctuation synchrony, we also examined the consistency of phase alignment between oscillations in the brains of the two participants in each pair (Imaginary Coherence). This metric represents much faster processes, on the order of tens of milliseconds, and is independent of the strength of neural activity. Mindful of the data quality disadvantage when conducting neuroscience research in a non-laboratory context, we took advantage of the large size of our dataset to adopt strict inclusion criteria. First, we decided to focus our analysis on only one recording site (1,568 people) for consistency purposes, which was further reduced to a group of 614 datasets that met a high data quality threshold. Furthermore, as discussed in the Introduction and described in the Methods, our synchrony metrics were chosen so that the effect of instantaneous brain signal co-fluctuations is minimized, circumventing contamination by environmental electrical noise present in the museum environment.

### Intrinsic motivation: relationship and personality traits predict inter-brain coupling

We first explored whether properties of pairs’ relationship predicted the average inter-brain coupling across the 10-minute experience (cf. Figure 1b). Cluster statistics revealed significant effects of both relationship duration (a trait measure) and social closeness (a state measure; (Aron, Aron, and Smollan 1992). People who knew each other longer showed stronger inter-brain coupling in the lower frequency ranges (Figure 4a, projected power correlations at 8 Hz; r(302) = .1776; p = .0019). Pairs who on average felt closer to each other also showed more inter-brain coupling with each other during the experience, in the beta-frequency range (Figure 4b; imaginary coherence at 21-22 Hz; r(307) = .1552, p = .005). Note that there was no difference in inter-brain coupling between strangers and pairs who already knew each other, in contrast to previous findings (Kinreich et al. 2017).

**Figure 4.**
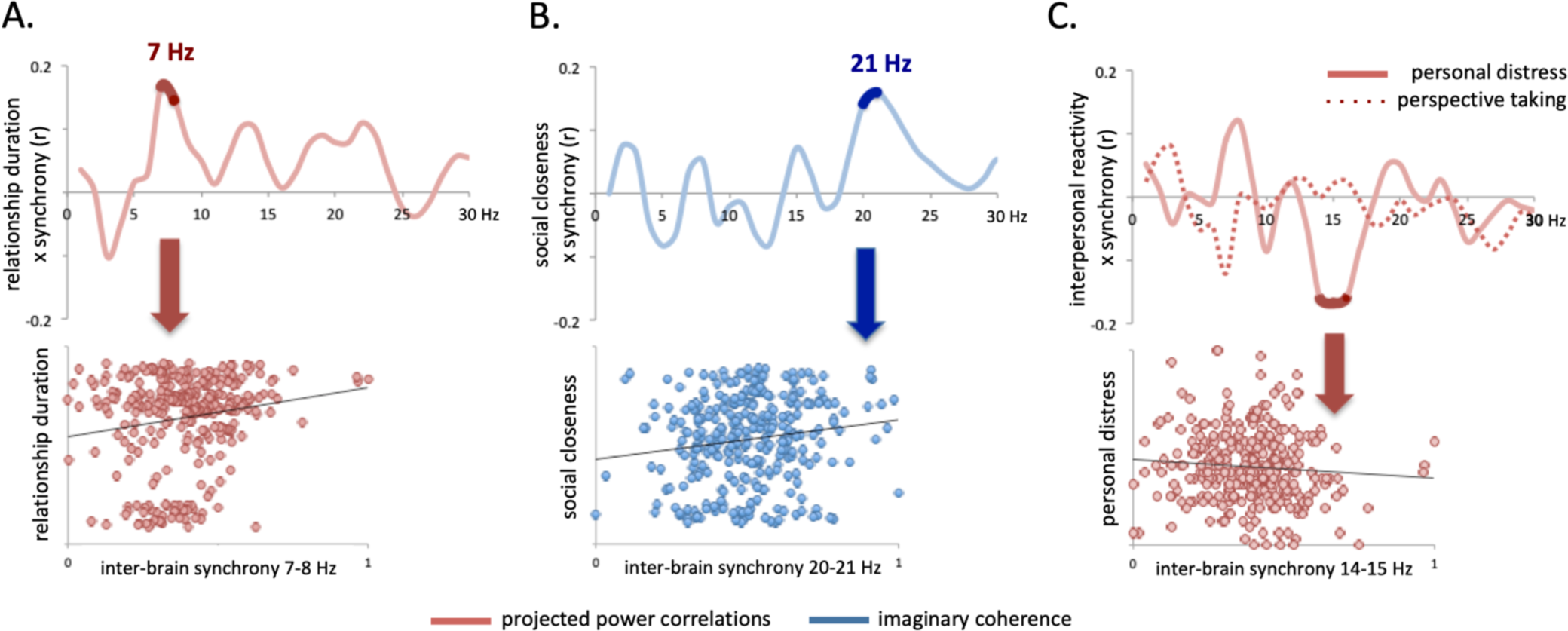
Inter-brain coupling is correlated with pairs’ relationship and with their affective personalities. Inter-brain coupling was significantly correlated with **(A)** relationship duration (projected power correlations at 7-8 Hz; r(302) = 0.1776; p = 0.0019) and **(B)** social closeness (imaginary coherence at 21-22 Hz; r(307) = 0.1552, p = 0.005). **(C)** A significant negative correlation was found between Personal Distress and inter-brain coupling (projected power correlations at 14-15 Hz; r(300) = 0.1757; p = 0.0023). No significant correlation between Perspective Taking and inter-brain coupling was found. Values are max-min normalized for presentation purposes and correlation plots show the average for each significant cluster.

Figure 4c shows the correlation plots for affective personality traits (Davis and Others 1980) and inter-brain coupling. Perspective Taking did not affect the average inter-brain coupling across the experience (dashed line), but Personal Distress did (solid line): At ∼15-16 Hz, there was a negative correlation between pairs’ average Personal Distress and their projected power correlations (*r*(300) = -0.1757, *p* = .0023), indicating that less emotionally self-oriented pairs (as measured through less Personal Distress), showed more inter-brain coupling overall.

### Pairs’ changes in focus predict inter-brain coupling

Next, we asked whether changes in pairs’ mental state, in particular focus level (Watson and Clark 1994) was associated with changes in inter-brain coupling (cf. Figure 1c). When comparing participants’ self-reported focus before and after the experience, we saw that 182 pairs showed a decrease in self-reported focus over time, while 84 pairs reported an increase in focus (all participant pairs Focus Pre vs. Focus Post: t(1, 452) = 6.8747, p < .0001; Focus Pre: M = 3.3392, SD = 0.6272; Focus Post: M = 3.1428, SD = 0.7224). For the group of pairs whose focus decreased, a smaller decrease was associated with a higher increase in projected power correlations at 6-7 Hz for the second compared to the first half of the experience (r(182) = 0.189, p = 0.0106). For the group who reported to be more focused after than before the experience, a higher increase in focus was associated with a higher increase in imaginary coherence (r(84) = 0.2975, p = 0.006). In other words: maintaining focus led to an increase in inter-brain coupling over time.

### Changes in low frequency inter-brain coupling are paired with changes in high frequency inter-brain coupling

Interestingly, both pairs’ self-reported focus and the nature of their relationship were correlated with projected power correlations at 7-8 Hz and imaginary coherence at 20-21 Hz respectively. We thus explored the relationship between the two inter-brain coupling measures at these two frequency ranges and found them to be coupled: changes in lower frequency projected power correlations at 7-8 Hz were positively correlated with changes in imaginary coherence at 20-21 Hz (Figure 5B: (r(73) = .2776, p = .0174).

**Figure 5.**
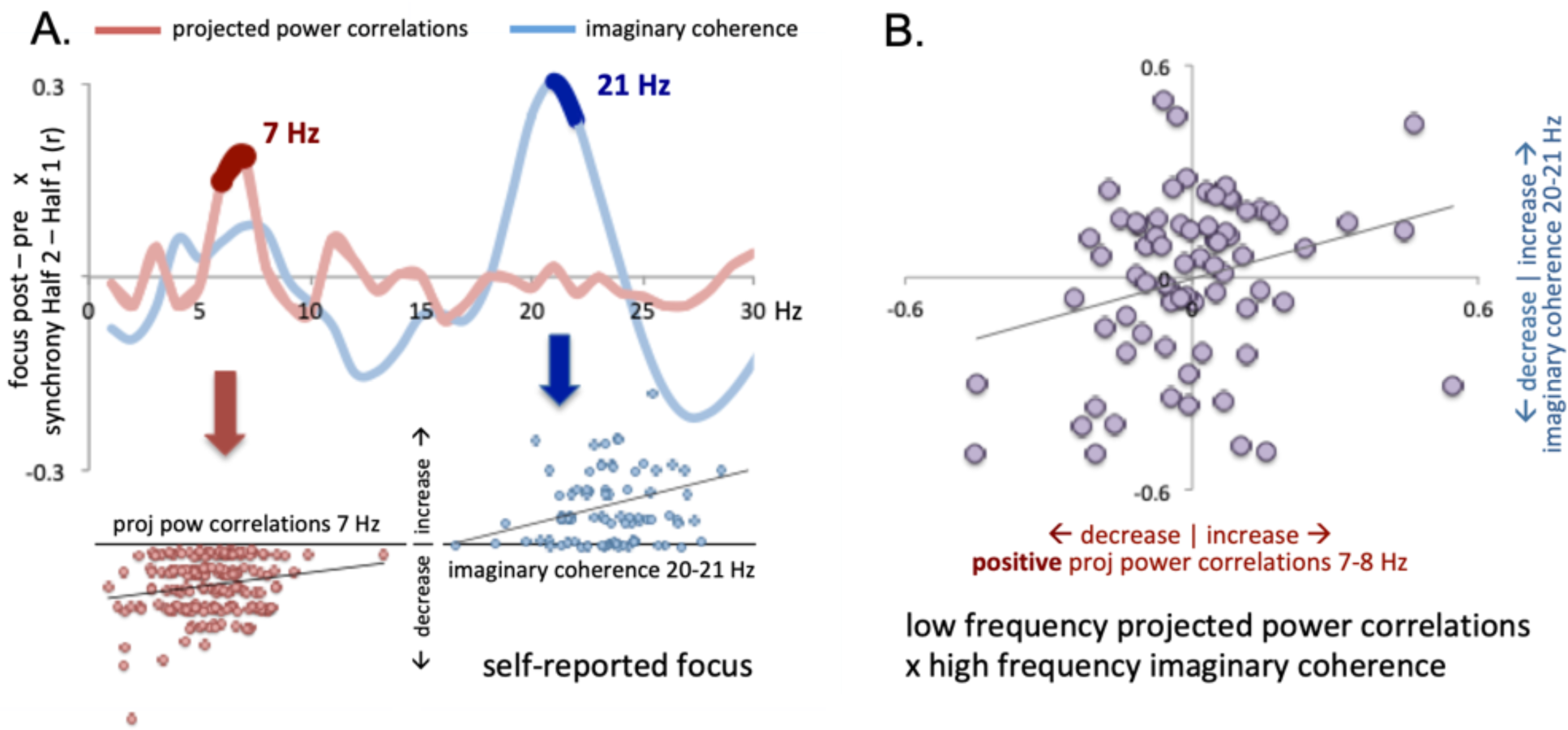
Inter-brain coupling is correlated with pairs’ focus level, and low frequency inter-brain coupling predicts high frequency inter-brain coupling (A-left) Pairs with a smaller decrease in focus and **(A-right)** pairs with a relatively higher increase in focus exhibited a relatively higher increase in inter-brain coupling in the second than the first half of the experience, in projected power correlations at 6-7 Hz (r(182) = 0.1889, p = 0.0106) and imaginary coherence at 20-21 Hz (r(84) = 0.2975, p = 0.006) respectively. **(B)** Changes in 7-8 Hz projected power correlations were positively associated with changes in imaginary coherence when comparing inter-brain coupling during the first half and the second half of the experience (r(73) = 0.2776, p = 0.0174). Pairs with negative projected power correlations are excluded. Values are max-min normalized for presentation purposes and correlation plots show the average for each significant cluster.

### Extrinsic motivation: inter-brain coupling is enhanced by explicit “synchrony” A/V

So far, we have shown that affective traits as well as mental states predict (changes in) inter-brain coupling during face-to-face interaction. We next asked whether pairs that were more motivated to connect also synchronized more (cf. Figure 1d). As described in the Methods, pairs from the BENAKI dataset were divided into two groups: one group was explicitly told that the visuals were derived from their correlated EEG signal (explicit synchrony A/V), while the other group did not (non-explicit synchrony A/V). We hypothesized that the feedback instructions would function as a motivational factor to remain focused on the other. Indeed, pairs in the explicit feedback group showed an increase in inter-brain coupling for the second vs. the first half of the experience (Figure 6; Half 1 vs. Half 2 projected power correlations at 18-21 Hz: t(1, 138) = 2.7049, p = 0.0077, M = 0.037, SD = 0.1619). One plausible explanation for this discrepancy is that the “explicit feedback” group was more motivated to maintain focused, or engaged with the other person throughout the 10-minute experience: the visual environment functioned as a constant reminder of the task, namely to connect to the person directly opposite.

**Figure 6.**
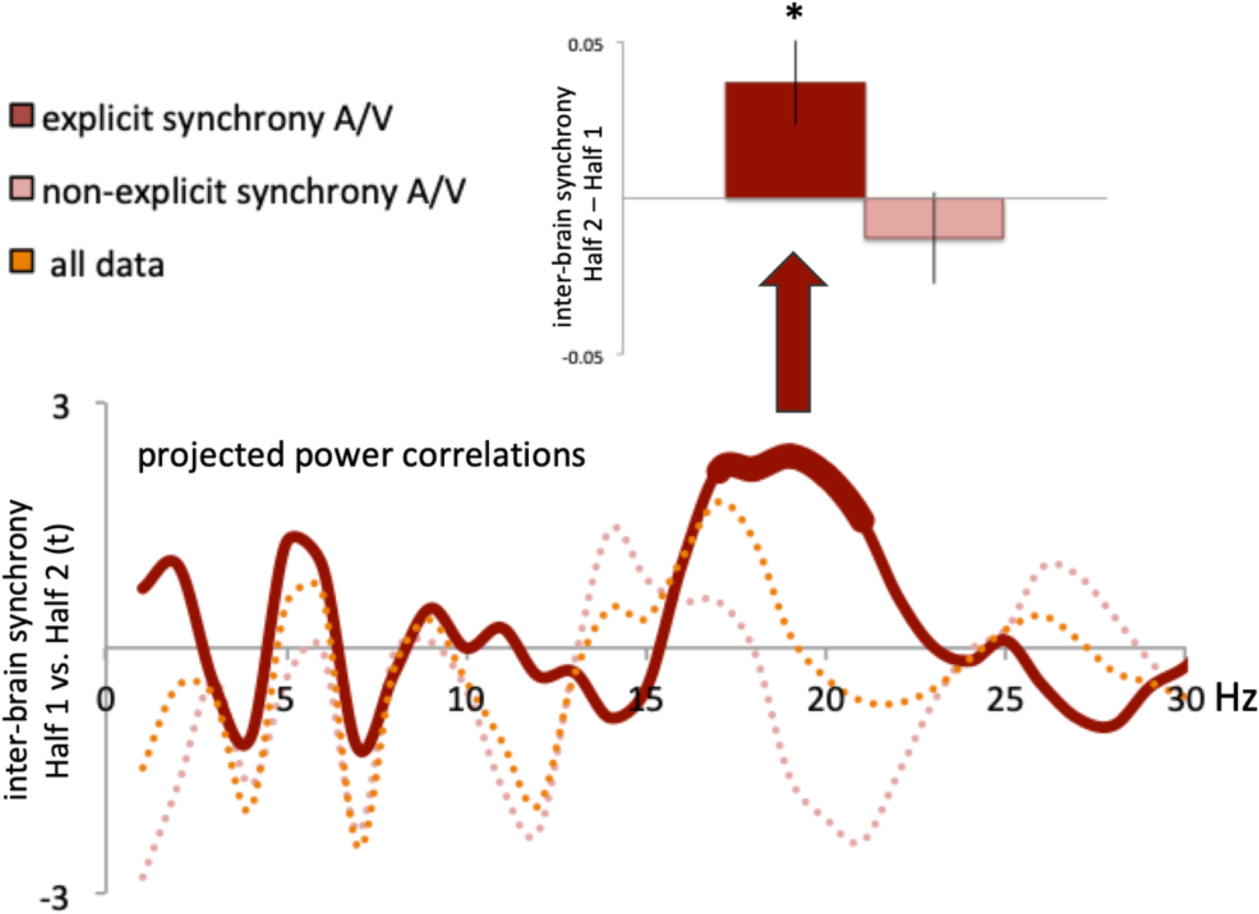
Extrinsic motivation to connect leads to an increase in inter-brain coupling. Pairs who received no explicit explanation of the relationship between the A/V environment and inter-brain correlations showed no significant changes in inter-brain coupling over time; Pairs in the explicit feedback group exhibited higher inter-brain coupling for the second vs. the first half of the experience (projected power correlations at 18-21 Hz: t(1, 138) = 2.7049, p = 0.0077, M = .037, SD = .1619). Values are max-min normalized for presentation purposes. Error bars reflect standard errors of the mean.

Indirect support for such a shared engagement account of inter-brain coupling comes from the following set of observations in our data. First, while the “no explicit feedback” group exhibited a decrease in focus (t(1, 209) = 3.1647, p = 0.0018, SD = .1499) for the post vs. pre questionnaire, the “explicit feedback” group showed no significant changes in focus. A second, related, observation is that self-reported focus did not predict changes in inter-brain coupling when restricting the analyses reported in Figure 5a to the “explicit feedback” group alone. Importantly, the relationship between inter-brain coupling and pairs’ relationship or personality traits was unaffected by the instructions that pairs received about the task and A/V environment (See supplemental Figure S1 for details). Thus, introducing extrinsic motivation to socially engage appeared to exclusively override the effect of self-reported focus on inter-brain coupling.

A final piece of evidence in support of the proposed role of shared engagement in inter-brain coupling comes from the finding that participants’ inter-brain coupling changes were not affected if they were presented with a sham A/V environment. Recall that half the pairs in the LOWLANDS dataset saw visuals that were randomly generated rather than being controlled by true inter-brain coupling values. Importantly, all pairs in the LOWLANDS dataset were explicit told that the A/V environment was related to their ongoing inter-brain coupling. As shown in Figure S2, in contrast to the BENAKI pairs, where only the “explicit feedback” group showed an increase in inter-brain coupling for the second compared to the first half of the experience, in the LOWLANDS dataset inter-brain coupling increased over time for all pairs irrespective of whether the inter-brain coupling environment was true or sham (projected power correlations at 16-18 Hz: t(1,55) = 3.2379, p = 0.002, SD = 0.0193; i.e., in the same frequency range as for the BENAKI data shown in Figure 6). This suggests that *believing* that the visuals were directly related to the success of the interaction motivated participants to remain socially engaged, irrespective of the actual relationship between inter-brain coupling and the visual environment.

Importantly, no other differences were observed between the two datasets: with the exception of social closeness, all findings reported in Figures 4 and 5 for the BENAKI dataset were replicated in the LOWLANDS dataset (See Figure S2 for details). A further minor difference was that the frequency ranges of the projected power correlations were a bit higher for the LOWLANDS dataset (∼10 Hz as opposed to 8 Hz). This could be due to age differences between the two groups: Alpha peak frequency is typically lower for older than for younger adults (e.g., (Duffy, McAnulty, and Albert 1993) and while no age information available for the BENAKI participants, the music festival Lowlands is known to attract a younger demographic than the Benaki art museum.

### Social behavior as an exogenous synchronizer

Finally, we asked whether the type of social behavior that pairs engaged in during the social interaction was predictive of their inter-brain coupling (Cf. Figure 1e; LOWLANDS dataset only). Participants listed a number of different strategies they used to try to synchronize with one another, which included “no strategy” (10.7% of pairs), and “stimulus entrainment” (focusing on the visuals: 37.5% of pairs), in addition to three main categories of social behavior: “eye contact” (71.4% of pairs), “joint action” (performing the same physical action such as smiling or having a conversation, playing hand games: 25% of pairs), and “joint thought” (thinking about the same object, event, or each other: 32 % of pairs). As can be seen in Figure 7a-b, “eye contact” and “joint action” were positively associated with inter-brain coupling (eye contact: projected power correlations at 9-11 Hz (r(56) = 0.3786, p = 0.0040) and 26-30 Hz (r (56) = 0.3509, p = 0.008); joint action: imaginary coherence at 18-21 Hz (r(56) = 0.3651, p = 0.0057), but “joint thought” was not predictive of inter-brain coupling (Figure 7c).

**Figure 7.**
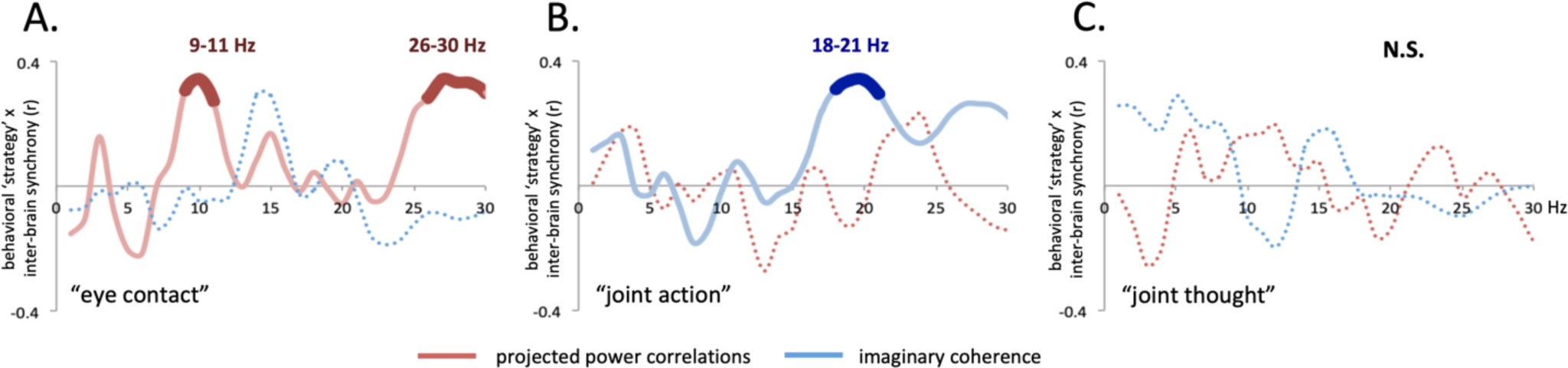
Synchronizing “strategies” predict inter-brain coupling in the Lowlands dataset. Using **(A)** “eye contact” or (**B)** “joint action” as strategies to connect was positively associated with inter-brain coupling (eye contact: projected power correlations at 9-11 Hz (r(56) = 0.3786, p = 0.0040) and 26-30 Hz (r (56) = 0.3509, p = 0.008); joint action: imaginary coherence at 18-21 Hz (r(56) = 0.3651, p = 0.0057), but (**C**) “joint thought” strategies were not. See main text for a description of the categories. Values are max-min normalized for presentation purposes.

## DATA AND RESULTS DISSEMINATION

All data is made publicly available at https://osf.io/hpgkt/ and the results are disseminated to participants and the general public via wp.nyu.edu/mutualwavemachine

## DISCUSSION

In an effort to explore the neural correlates of real-world social behavior, we present data acquired in a novel context departing from laboratory-constrained cognitive neuroscience. Building on recent technical and analytic advances in recording neural data from groups in natural settings, we extended such data acquisition approaches in a new direction, recording EEG from a very large number of people, recruited from the general public, as they engaged in naturalistic face-to-face interaction. Specifically, we created an interactive neurofeedback art experience, the Mutual Wave Machine, which allowed us to ask how pairs’ relationship, personality traits, mental states, and social behavior predicted inter-brain coupling during face-to-face interactions, extending ongoing laboratory research on neuronal oscillations and their role in perception and cognition, as well as previous EEG hyperscanning studies ((F. Babiloni et al. 2007; Dumas et al. 2010; Pérez et al. 2018; Kinreich et al. 2017; Sänger, Müller, and Lindenberger 2012); and many others). We employed a ‘crowdsourcing’ neuroscience approach, recruiting museum visitors and festival goers to help us explore our research questions. For example, we asked participants to indicate what kind of behavior *they* thought had helped them synchronize their brain activity.

In a subgroup of 726 participants whose data survived rigorous criteria licensing further analyses, we found that inter-brain coupling was positively related to pairs’ social closeness, personality traits, focus level, and motivation to connect (Figure 4A/B, Figure 4C, Figure 5A, and Figure 6 respectively). Further, modulations in alpha synchrony (projected power correlations) co-varied with changes in beta coherence (at 21-22 Hz), suggestive of a relationship between lower and higher frequency inter-brain coupling (Figure 5B).

### Social closeness, personality traits, and focus as predictors of synchrony

As reviewed in the Introduction, our findings corroborate EEG hyperscanning studies that have reported relationships between inter-brain coupling on the one hand, and social closeness, personality traits, and individual focus on the other (for a recent review see e.g., (Czeszumski et al. 2020), and extend these to more naturalistic social interactions than typically employed. However, it is worth noting that studies vary in the metrics used, in terms of both the computation of inter-brain coupling ((Ayrolles et al., in revision) and the assessment of e.g., personality traits. This may lead to a false impression of consistency between findings. To illustrate, we find that Personal Distress but not Perspective Taking predicts inter-brain coupling, corroborating our own prior findings in classroom interactions (e.g., (Dikker et al. 2017). However, other research (Goldstein et al. 2018) has instead found that Perspective Taking but not Personal Distress predicts inter-brain coupling, quantified in their case using “cCorr” (circular correlation coefficient; (Burgess 2013). This is just one illustration of how “empathy” and “synchrony” may actually be referring to completely different constructs depending on the study, and that the hyperscanning field has yet to reach consensus on the extent to which these differences between studies are cognitively meaningful.

### Shared attention and engagement

In line with previous work, our findings invite an interpretation based on shared attention, or shared engagement (Dikker et al. 2017; Ki, Kelly, and Parra 2016; Cohen, Henin, and Parra 2017), illustrated in Figure 8.

**Figure 8.**
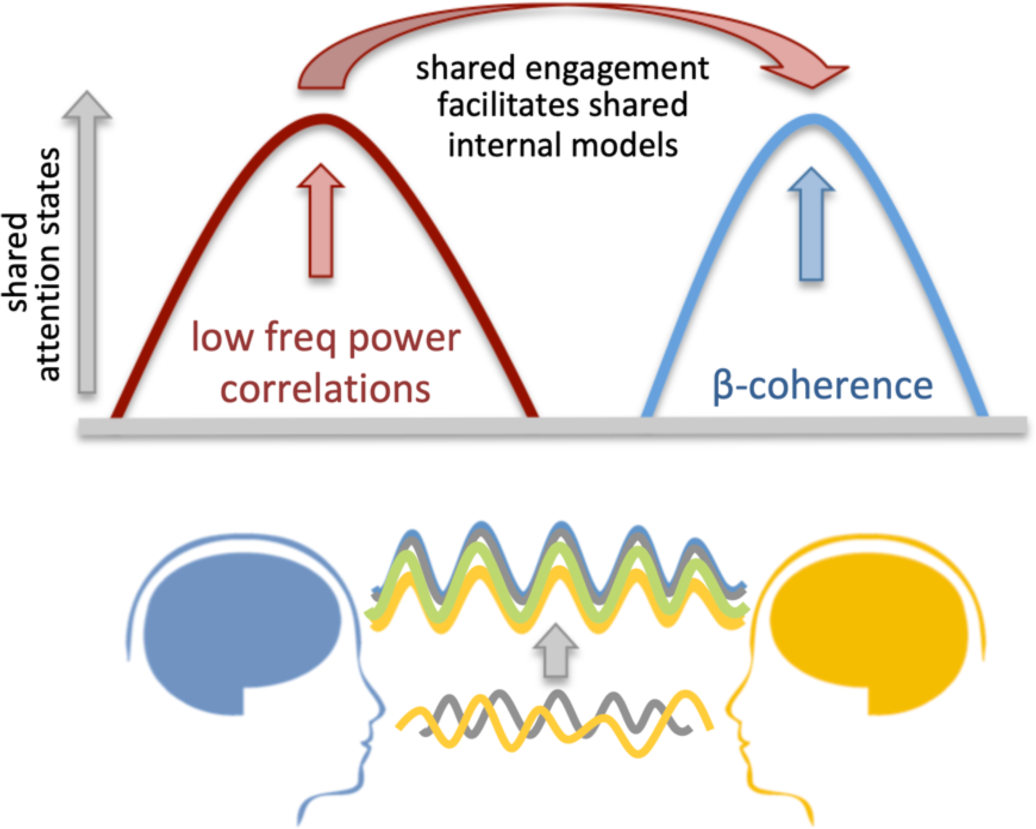
Shared engagement facilitates the formation of shared internal models. A schematic model showing how two people who have more instances in shared attention (‘moments of meeting’) can be measured as similar low frequency power changes, which in turn enables the tuning of shared internal models.

Projected power correlations, by hypothesis, would capture whether pairs show concurrent changes in attentive states, with positive correlations indicating that they are (in)attentive at the same time and low correlations or negative correlations indicating that they do not often share the same attentional state during the experience. ‘Synchrony’, thus, does not imply that pairs maintain a high focus level throughout the experience, just that their in-and-out of attention states co-fluctuate. Pairs who are more often in an attentive state together (similar alpha power changes) are more likely to simultaneously pay attention to each other’s actions or other cues from the surrounding environment, resulting in more similar neural representations or predictions (Arnal and Giraud 2012). In other words, we hypothesize that the findings shown in Figure 5B invite an interpretation where shared attention states facilitate interpersonal neural synchrony at the oscillatory phase level (beta synchrony).

Inter-brain synchrony in the beta band as a function of (social) attention converges with research showing that joint action is supported by oscillatory activity in the beta frequency range. Beta/mu rhythms, typically measured over sensorimotor areas (Pineda 2005; Hari 2006), have been associated with attention (Anderson and Ding 2011), motor control and motor simulation (Pfurtscheller and Lopes da Silva 1999), as well as prediction of another person’s actions (Sebanz, Bekkering, and Knoblich 2006). Changes in beta/mu are observed both when people perform an action and when they watch someone perform a similar action (Nishitani and Hari 2000). Crucially, beta activity during action perception varies as a function of social evaluation (Koelewijn et al. 2008) as well as social traits such as empathic concern (Perry, Troje, and Bentin 2010) and, in line with our findings, Personal Distress (Saarela et al. 2007; Yang et al. 2009). Our work also links to previous EEG studies comparing neural oscillations between people during interpersonal coordination tasks (Fabio Babiloni and Astolfi 2014; Dumas et al. 2010; Sänger, Müller, and Lindenberger 2012; Szymanski et al. 2017). For example, recent work by Novembre and colleagues (Novembre et al. 2017) showed that dual in-phase 20 Hz brain stimulation enhanced interpersonal movement synchrony between two participants performing a finger-tapping task. We expand on these findings by showing that brain-to-brain synchrony in the beta frequency range is sensitive to interpersonal factors such as affective personality traits during real-world face-to-face social interaction, which can further be linked to research that has associated beta oscillations with endogenous content representations and expectations (Arnal and Giraud 2012; Spitzer and Haegens 2017).

In sum, we propose an account wherein shared attention (Kang and Wheatley 2017; Leong, Byrne, and Clackson 2017; Dikker et al. 2017; Ki, Kelly, and Parra 2016) is measured via an increase in projected power correlations in the alpha frequency range. Concretely: only when both members of the dyad are paying close attention to the interaction, which has been shown in numerous studies to be correlated with low alpha power, will they be able to establish shared (motor, perceptual, cognitive) representations, which we hypothesize is reflected in an increase in imaginary coherence at 20 Hz. It is important to emphasize that our results do not directly test this dissociation, so this interpretation remains hypothetical here and should be tested in future studies where attention and joint action are manipulated independently.

### Neurofeedback validity

Typically, neurofeedback modules used in BCI setups are validated by comparing online output to offline analyses. In our case, a different inter-brain connectivity metric was used for the online neurofeedback setup than for subsequent offline analysis, deviating from this common practice. This is in part due to limitations mentioned above: We did not implement online artifact rejection (e.g., Mullen et al. 2015) and the narrow-band analysis approach increased the likelihood of inflating correlation values. In addition, and perhaps most importantly, in contrast to other EEG signatures that are commonly implemented in BCIs (such as the P3 ERP component; e.g., Fazel-Rezai et al. 2012), the correlational approach used here has not been extensively validated in laboratory experiments (e.g., Czeszumski et al. 2020). In ongoing work, we are evaluating and benchmarking inter-subject connectivity metrics (Ayrolles et al., in revision) which are then implemented into a graphical user interface and systematically validated for inter-brain BCI purposes (https://github.com/rhythmsofrelating).

### On conducting “neuroscience in the wild”

Carrying out “crowdsourcing” neuroscience research outside of the laboratory comes with many benefits, but also a number of challenges. First, it is near-impossible to obtain full experimental control in public spaces, and this project was especially challenging in this regard. For example, the LOWLANDS dataset was collected during a music festival, which required extra care with respect to noise contamination from surrounding events. Further, due to the sheer number of participants as well as other logistical and privacy-related considerations, we were unable to keep a close record of participant behavior during the interaction. As a result, we had to rely on participant self-report (see Figure 7), which provided us with information to assess the brain-behavior relationship during the social interactions, but of course this information was incomplete.

Another challenge was the hybrid art/science/tech nature of the Mutual Wave Machine. While participants took their roles seriously in both BENAKI and LOWLANDS, for a few other sites listed in Table S1, visitors did not treat the experience as a scientific experiment but rather as a curiosity (e.g., taking selfies instead of interacting with each other). People were also often waiting in line to participate, sometimes even getting impatient, jeopardizing the setup and reliability of their questionnaire responses. At these sites, the experience was shortened to 6 minutes or less. For these and other reasons, we only analyzed datasets that were collected during multiple days and where 20-minute timeslots were assigned with 5-to 10-minute buffers on each side. At BENAKI, participants further self-selected by signing up in advance via an online portal.

Another challenge relates to equipment. The EEG devices used here (EMOTIV) were very suitable for our purposes because they are sturdy, fast to apply, easy to handle, and affordable. However, data quality may be lower compared to laboratory-grade equipment (Krigolson et al. 2017). As discussed in the Methods, we took various steps to ensure that our data met rigorous standards despite these limitations.

On the flipside, the benefits of conducting neuroscience research outside of traditional laboratory environments are clear. First, using a citizen science approach affords researchers the opportunity to collect data from large numbers of people with a more varied demographic profile than the typical participant population of laboratory neuroscience research. Further, actively involving the general public in research has a number of benefits beyond constituting a rich opportunity for neuroscience outreach and education: While it is common to view interactions between scientists and the general public as merely unilateral (scientists educate the public about their work), we would like to argue that interactions with artists, educators, and the general public can inform scientific inquiry in a fruitful way: non-specialists may force scientists to remain aware of any translational value of their work to everyday practice, challenge methodological approaches that are taken for granted, and inspire research questions that may inform laboratory research. To close with an emphasis on the latter point: while the field may be ‘ready’ for real-world neuroscience (Matusz et al. 2019a), in our opinion it will flourish only if paired with rigorous laboratory-based work and solid, careful methodology.

## CONCLUSION

A large group of museum and festival visitors engaged in dynamic face-to-face interactions while their brain activity was recorded using EEG. This setup made it possible to explore the limits and opportunities afforded by conducting human social neuroscience research outside of the traditional laboratory context. Drawing on our two most comprehensive datasets to date, we were able to evaluate how intra- and interpersonal factors predict the extent to which brain activity becomes synchronized between people during face-to-face interaction. Pairwise synchronized brain activity was related to people’s relationship, affective personality traits, mental states, as well as their motivation and strategy to connect to the other person. We propose an account for brain-to-brain synchrony in which shared engagement provides a vehicle for synchronous brain activity (measured in the alpha and beta frequencies, respectively), and joint action is used to mutually adapt neural and behavioral representations. Taken together, we demonstrate that an unconventional, ‘crowdsourcing neuroscience’ approach can provide valuable insights into the brain basis of dynamic real-world social behavior.

## Acknowledgements

This research was supported by the Netherlands Organization for Scientific Research Awards 275-89-018 and 406.18.GO.024. The Mutual Wave Machine was made possible with support by Creative Industries Fund NL, TodaysArt, Marina Abramovic Institute, de Hersenstichting, Lowlands Science, Utrecht University, and NEON.

## Design, tech & production

Peter Burr, Danielle Boelling, Diederik Schoorl, Jean Jacques Warmerdam, Matthew Patterson Curry, Pandelis Diamantides;

## Data collection and management

Annita Apostolaki, Dana Bevilacqua, Shaista Dhanesar, Imke Kruitwagen, Eletta Daemen, Orsa Rebouskou, Stella Papazisi, Aspa Papazisi, Karlijn Blommers, Sascha Couvee, Ella Bosch, Jorik Geutjes, Chris van Run.

