## Supplemental Materials for "Crowdsourcing neuroscience: inter-brain coupling during face-to-face interactions outside the laboratory"

| **`Mutual Wave Machine Locations** | **Year** | **Number of Participants** |
|---|---|---|
| TodaysArt Festival - The Hague | 2013 | 108 |
| Eye Film Institute - Amsterdam | 2013 | 56 |
| Nemo Science Museum - Amsterdam | 2014 | 64 |
| Silicon Valley Contemporary - San Jose, CA | 2014 | 208 |
| Lexus Hybrid Art - Moscow | 2014 | ~ 600 |
| **Lowlands Festival (LOWLANDS) - Netherlands** | **2015** | **230** |
| Benaki Museum **(BENAKI) - Athens** | **2016** | **1568** |
| Innovation Expo - Amsterdam | 2016 | ~ 50 |
| Tivoli/Vredenburg - Netherlands | 2016 | ~ 100 |
| 3LD Art and Technology Center - NYC | 2016 | ~ 80 |
| FORMS Festival - Toronto | 2016 | 66 |
| Pioneer Works - NYC | 2017 | 52 |
| OPUS 1 Festival - Merriweather, MD | 2017 | ~ 150 |
| Espacio Telefónica - Madrid, Spain | 2019 | ~ 1470 |
| **Total Nr. of Participants** | | **~ 4800** |

**Table S1. Mutual Wave Machine Locations**
List of locations/venues where the Mutual Wave Machine has been set up since 2013, with number of participants per location.

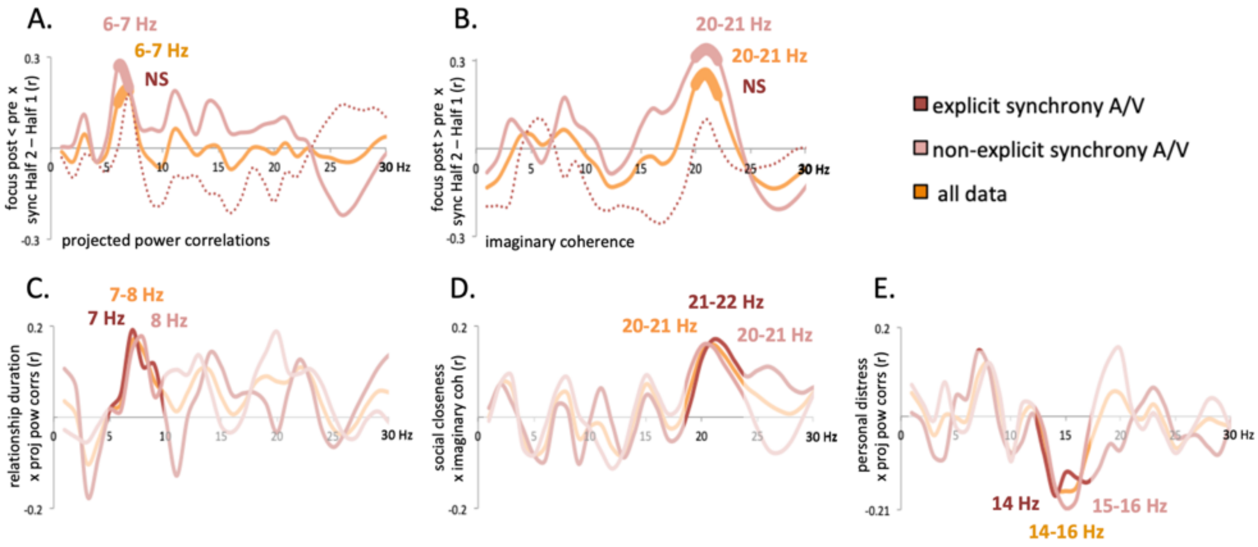


**Figure S1. Inter-brain synchrony results for BENAKI, divided by group**

**(A-B)** Changes in self-reported focus between the pre and post questionnaire (focus pre < focus post) were positively correlated with changes in inter-brain synchrony between the 1st and 2nd half of the experience (Half 2 – Half 1) for all data (orange line, replotting Figure 5a), and the "no explicit feedback" pairs (pink line, A. projected power correlations at 6-7 Hz: $r(101) = 0.2684$, $p = 0.0066$; B. Imaginary coherence at 20-21 Hz: $r(67) = 0.3340$, $p = 0.0049$), but not for pairs who received explicit instructions about the relationship between synchrony and the A/V environment ("explicit feedback" group, red dotted line). Importantly, inter-brain synchrony correlations with **(C)** relationship duration (all data: orange line, replotting Figure 4a; "explicit feedback" pairs: red line, projected power correlations at 7-8 Hz: $r(138) = 0.1910$, $p = 0.0248$; "no explicit feedback" pairs: projected power correlations at 7-8 Hz: $r(164) = 0.1749$, $p = 0.0251$) **(D)** social closeness (all data: orange line, replotting Figure 4b, orange line; "explicit feedback" pairs: red line, imaginary coherence at 20-21 Hz: $r(138) = 0.1697$, $p = 0.0458$; "no explicit feedback" pairs: pink line, imaginary coherence at 20-21 Hz: $r(168) = 0.1548$, $p = 0.0457$) and **(E)** personal distress (all data: orange line, replotting Figure 4c; "explicit feedback" pairs: red line, projected power correlations at 14-15 Hz: $r(136) = -0.1784$, $p = 0.0377$; "no explicit feedback" pairs: projected power correlations at 15-16 Hz: $r(164) = -0.2068$, $p = 0.0079$) were not affected by whether or not participants were informed of the mapping of inter-brain synchrony to the A/V environment. Values are max-min normalized for presentation purposes.

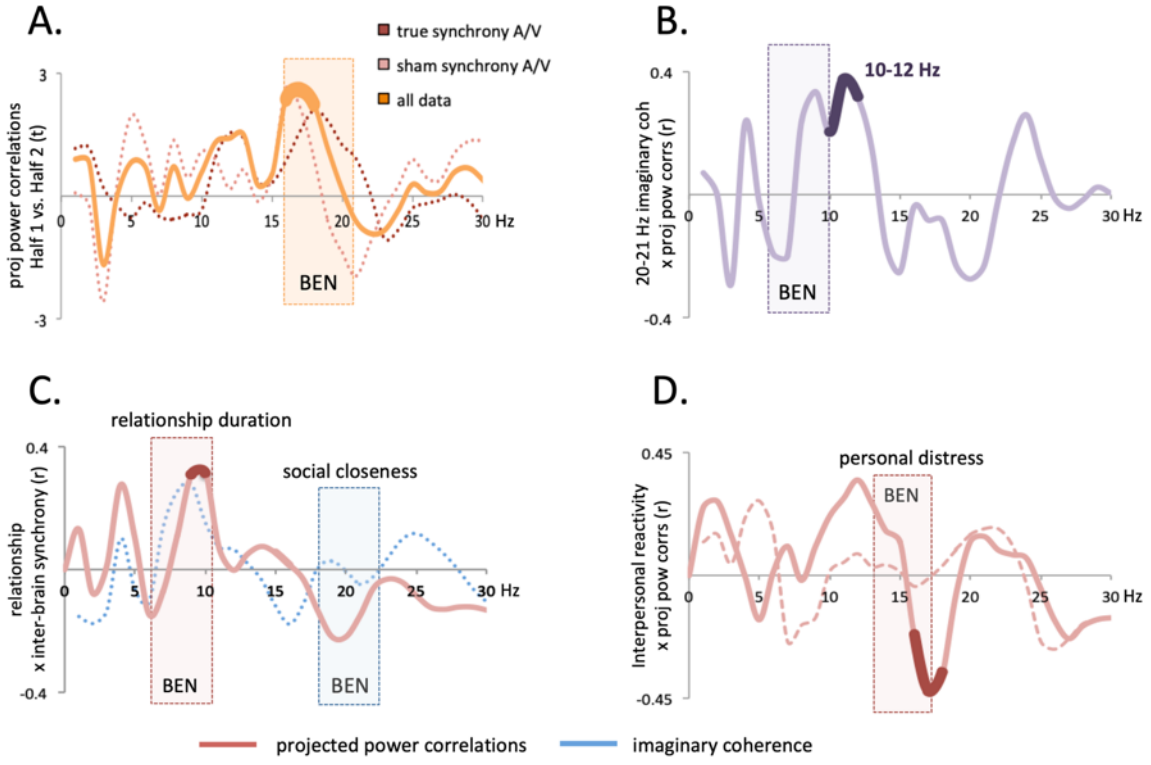


**Figure S2. Replicating inter-brain synchrony findings in the LOWLANDS dataset**

Inter-brain synchrony values for the LOWLANDS dataset are shown in relation to the same analyses in the BENAKI dataset. The latter are indicated with shaded areas superimposed on each panel. **(A)** Inter-brain synchrony increased over time in all data, independently of whether the A/V environment was dictated by random or true inter-brain synchrony values (all data: orange line, Half 1 vs. Half 2 projected power correlations at 16-18 Hz: $t(1,55) = 3.2379$, $p = 0.002$, $SD = 0.0193$; see Figure 2 and compare to Figure 6). **(B)** Changes over time in imaginary coherence at 21-22 Hz were positively correlated with changes in projected power correlations at 10-12 Hz ($r(56) = 0.4779$, $p = 0.001$, compare to Figure 5b). **(C)** In contrast to the BENAKI dataset, relationship length but not social closeness was correlated with inter-brain synchrony in the LOWLANDS dataset (relationship length: red line, projected power correlations at 9-10 Hz: $r(56) = 0.3107$, $p = 0.0198$, compare to Figures 4a and 4b). **(D)** Just like in the BENAKI dataset, personal distress (solid line) but not perspective taking (dotted line) was correlated with inter-brain synchrony in the LOWLANDS dataset (personal distress:, projected power correlations at 17-18 Hz: $r(56) = -0.4074$, $p = 0.0025$, compare to Figure 4c).

**S3. Text for general public website**

Non-hyperlinked (internal) links are indicated with < parentheses > and +/- indicates expandable sections

### THE MUTUAL WAVE MACHINE

Welcome! On this page you can explore the research findings from the Mutual Wave Machine, a “crowd-sourcing” neuroscience experiment: scientists asked the general audience to help them understand what happens in our brains when we interact face-to-face with another person. If you were one of those participants: **THANK YOU!**

**INDEX**

**<WHAT IS THE MUTUAL WAVE MACHINE?> <WHO PARTICIPATED?>**
**<HOW DOES IT WORK?> <WHAT DID WE FIND?> <CREDITS AND REFERENCES>**
**<MORE INFO ABOUT THE METHODS>**

#### WHAT IS THE MUTUAL WAVE MACHNE?

**What does it mean to be ‘on the same wavelength’ with another person? When you feel connected to another person, are your brainwaves literally ‘in sync’?**

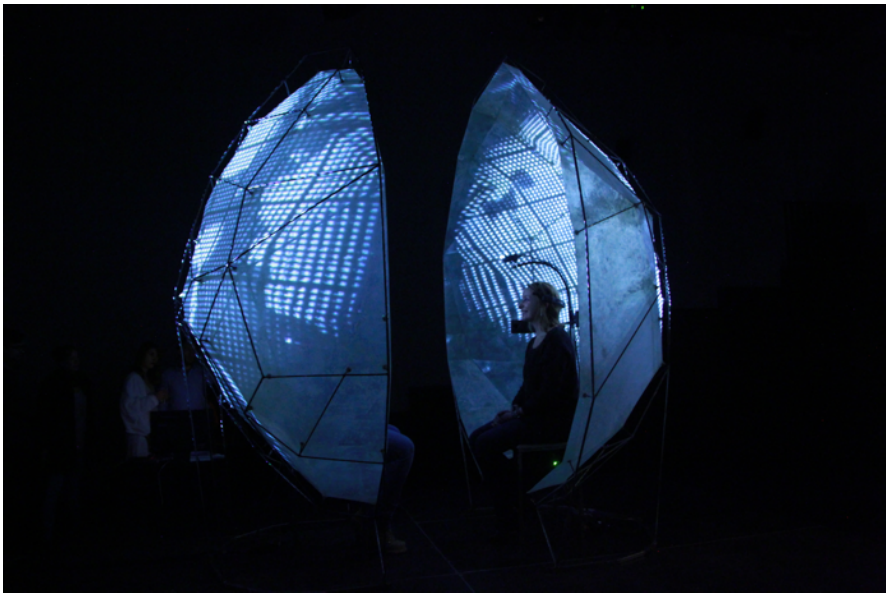

< embedded video showing the Mutual Wave Machine https://vimeo.com/96287858 >

The Mutual Wave Machine is a brain experiment meets interactive art installation. We use a **crowd-sourcing neuroscience approach to find out when our brainwaves synchronize with each other during dynamic social interactions**. In other words: researchers collaborate with the general public to explore what makes people 'click'.

Thousands of museum and festival visitors have worn EEG headsets to record their brainwaves while they sat inside the Mutual Wave Machine and interacted face-to-face with a stranger or loved one for 10 minutes. In the meantime, they were immersed in a real-time reflection of their ongoing brainwave synchrony (or, more accurately: correlations of their EEG signals, as explained below).

In 2016, The Mutual Wave Machine won the Art of Neuroscience Award.

Interested in exploring the data yourself? All the datasets will live here

<expandable/collapsible section>

**+/- BACKGROUND: BRAINS IN SYNC DURING REAL-WORLD SOCIAL INTERACTIONS**

**From studying the brain in isolation to studying the brain in conversation**

Most neuroscientists study brains in isolation, that is to say: people come to the laboratory and watch images or listen to sounds while their brain activity is recorded. This way of studying human perception and action, in a very controlled and detailed manner, has led to many important brain discoveries. But, as you can imagine, there are also some limitations. To start, it is not immediately obvious that laboratory tasks (e.g., clicking on a button as soon as possible when you see a green line on the screen) are representative of the real-world perceptual problems that our brains have to solve. Second, people who come to the lab are often university students, which may not be a fully representative demographic. Finally, if you are interested in how our brains support our social interactions with other humans, this approach is lacking in that people rarely interact with another person during laboratory studies.

The Mutual Wave Machine takes a new data collection approach to try to tackle these limitations in laboratory research: we measure the brain activity of people of all ages and backgrounds as they interact with each other naturally.

Specifically, we asked **when brain activity becomes synchronized between people.**

It has been known that synchrony between people is an important phenomenon in social cohesion. Take dancing for example, or even walking down the street: we automatically synchronize our moves. Recently, researchers have begun to study such synchrony at the brain level: taking a 'hyperscanning' approach, neuroscientists record brain activity from participants during social interaction and compare brain activity between them. So far, factors that have predicted inter-brain synchrony (= similarities in the brain responses of people) include: (i) listening to or watching the same stimulus (ii) social coordination, like conversation or joint action (iii) social intentions, like cooperation vs. competition and (iv) social closeness, for example how much students like their classmates.

If you are interested in reading about all of our synchrony-based art/science work, see this review

#### WHO PARTICIPATED IN THE MUTUAL WAVE MACHINE?

Nearly 5000 people have participated in the Mutual Wave Machine thus far, and thousands more have seen the experiment in action. Scroll over the map and explore the links to get basic information about each exhibition site. <See Table S1>

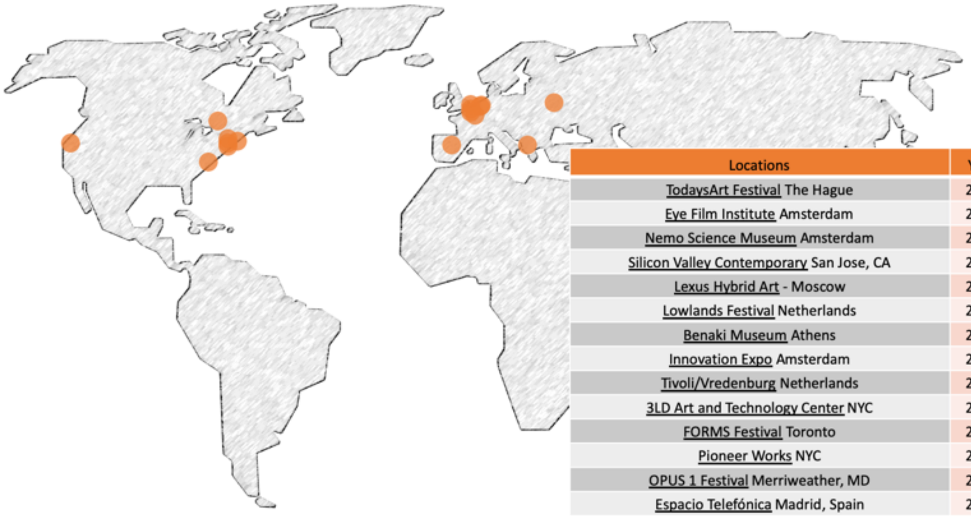

| Locations | Year | N |
|---|---|---|
| TodaysArt Festival The Hague | 2013 | 108 |
| Eye Film Institute Amsterdam | 2013 | 56 |
| Nemo Science Museum Amsterdam | 2014 | 64 |
| Silicon Valley Contemporary San Jose, CA | 2014 | 208 |
| Lexus Hybrid Art - Moscow | 2014 | ~ 600 |
| Lowlands Festival Netherlands | 2015 | 230 |
| Benaki Museum Athens | 2016 | 1568 |
| Innovation Expo Amsterdam | 2016 | ~ 50 |
| Tivoli/Vredenburg Netherlands | 2016 | ~ 100 |
| 3LD Art and Technology Center NYC | 2016 | ~ 80 |
| FORMS Festival Toronto | 2016 | 66 |
| Pioneer Works NYC | 2017 | 52 |
| OPUS 1 Festival Merriweather, MD | 2017 | ~ 150 |
| Espacio Telefónica Madrid, Spain | 2019 | ~ 1470 |

#### HOW DOES THE MUTUAL WAVE MACHINE WORK?

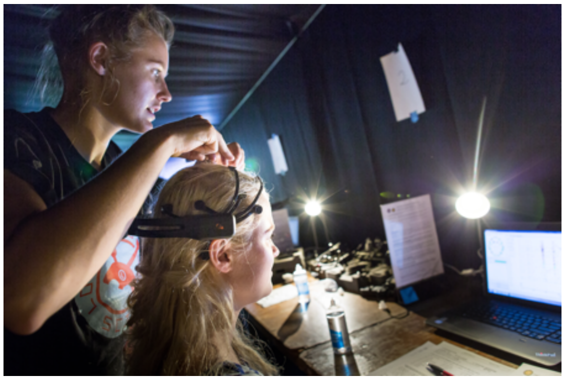

**Step 1.** If you volunteer to participate, you are first matched with a 'brainwave partner'. This can be someone you know or a complete stranger. You are each fitted with an EEG headset and you are shown your own <brainwaves> on a tablet or computer. You can clearly see why researchers usually ask people to sit still: the signal goes all over the place when you move (see <EEG> for more on this).

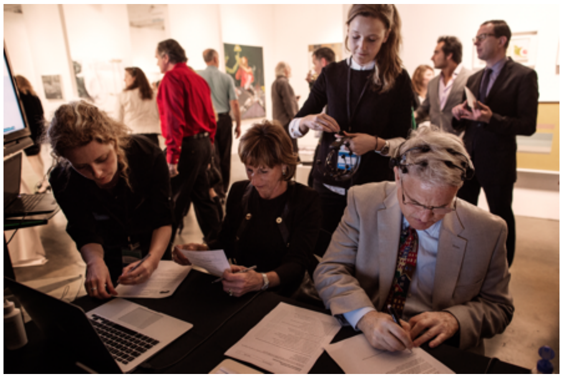

**Step 2.** Right before entering the Mutual Wave Machine, you each fill out a short questionnaire about your relationship, how you tend to behave in specific social situations, and your current mood.

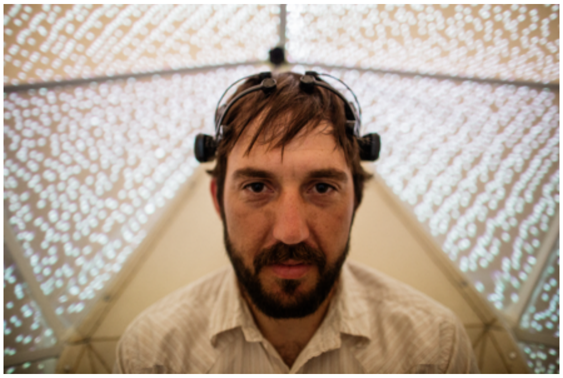

**Step 3.** Then you are led into the Mutual Wave Machine. For 7-10 minutes, you will sit opposite your brainwave partner and you will see and hear direct reflections of the correlations of your EEG signal (a proxy for "brainwave synchrony"): more light means more synchrony, and less light means less synchrony. See below for more information about this <neurofeedback>

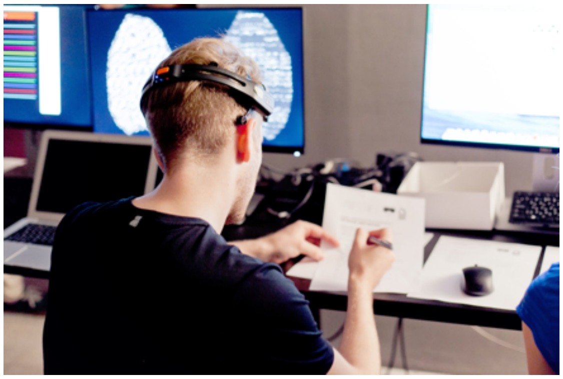

**Step 4.** After the experience, you will again fill out a short questionnaire.

#### BRAIN SYNCHRONY AND NEUROFEEDBACK

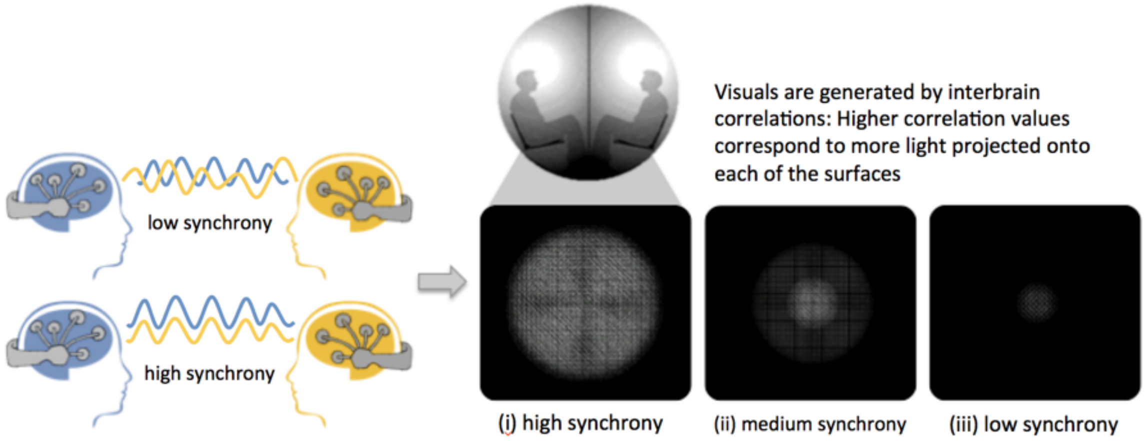


<Embedded video: https://vimeo.com/346434701>

The audio-visual environment displays, in real-time, how correlated the EEG signal is between two participants. As illustrated in the video and graphic, higher correlation values correspond to more light, and vice versa.

If you're interested in the exact calculations, see the detailed <Methods> below. See the detailed results for what effect this environment had on people's interactions.

#### WHAT DID WE FIND?

<expandable sections collapsed>

**+ DOES YOUR SOCIAL BOND AFFECT BRAINWAVE SYNCHRONY?**

**+ DOES *HOW* YOU INTERACT AFFECT YOUR BRAINWAVE SYNCHRONY?**

**+ DO PERSONALITY AND MENTAL STATE PREDICT BRAINWAVE SYNCHRONY?**

**+ DOES INTERACTING WITH EACH OTHER AFFECT YOUR MOOD AND SOCIAL BOND?**

**+ DOES THE "NEUROFEEDBACK" ENVIRONMENT AFFECT BRAINWAVE SYNCHRONY?**

<expandable sections expanded>

#### +/- DOES YOUR SOCIAL BOND AFFECT BRAINWAVE SYNCHRONY?

**Our main research question was whether feeling in sync with someone also syncs up your brainwaves with that person. So, we asked all participants to indicate how close they felt to each other (Social Closeness) and how long they had known each other (their Relationship Duration)**

In the graph below, each dot represents a pair of participants. If you participated in the Mutual Wave Machine, you and your "brainwave partner" may be one of those dots. The dots on the bottom of the graph represent pairs who felt less close to each other. Dots on the top of the graph represents pairs who felt very close to each other. Dots on the left of the graph represent pairs with low levels brain synchrony, pairs on the right of the graph represent pairs with relatively high levels of brain synchrony.

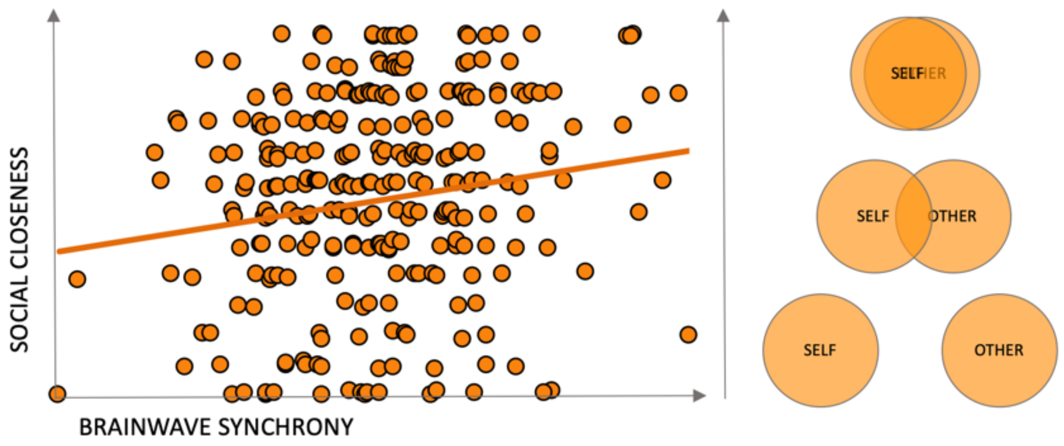


Overall, pairs who reported higher mutual social closeness showed more brain synchrony than pairs who did not feel close at all. However, as you can see, not every pair with high social closeness exhibited more brain synchrony than pairs who didn't know each other.

<video demonstrating interactive feature: https://vimeo.com/349465506

*If you were a participant, drag the "self" and "other" circles across each other or away from each other to know which set of dots may represent you and your brainwave partner. If you did not participate, you can imagine yourself in a position where you are interacting with someone you know.>*

Be sure to check out Carolyn Parkinson and Thalia Weatley's fMRI work showing that when you watch video clips, your brain responses are similar to those close to you in your social network.

#### +/- DOES *HOW* YOU INTERACT AFFECT YOUR BRAINWAVE SYNCHRONY?

Participants used different strategies to sync up their brainwaves. Many people tried: (1) doing something together (2) thinking about the same thing, or (3) eye contact. Which of these strategies do you think made brain synchrony go up?

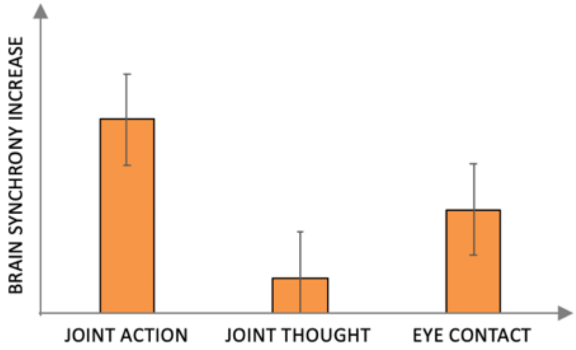


As you can see in the graph, doing something together (joint action) and eye contact are both associated with an increase in synchrony over time. Thinking about the same thing, however, did not appear to increase synchrony.

(1) **Doing something together (also referred to as "joint action")**: This includes singing a song, having a conversation, smiling, holding or moving hands together, etc. The first EEG 'hyperscanning' study that was published, by Guillaume Dumas, already showed that if you do something together, your brainwaves sync up. Later studies by Dumas and others have shown that conversation also synchronizes your brainwaves.

(2) **Thinking about the same thing**: This includes strategies like retrieving the same memories ("remember when we got stuck in the rain that one time?"), thinking about the same object or experience ("let's think about ice cream together!"), or thinking about favorite bands or movies. We didn't find any evidence that this kind of 'common grounding' puts people's brains on the same wavelength. One possible explanation is that EEG isn't precise enough to capture such joint thought, perhaps because the signal changes too fast. fMRI studies, which measure brain activity at a much slower pace, have shown that our brains respond similarly if we recall the same past experience or think about the same object.

(3) **Eye contact**: Many pairs used eye contact as a strategy to connect to each other, and we saw that this increased the synchronization of their gamma brain waves, in line with recent findings from other research groups. Some studies looking at brain synchrony between mothers and their babies have also shown that the It is not surprising that eye contact increases synchrony: eye contact, and gaze more generally, is a very strong social tool that is pivotal in human relationships. This finding also links to our work Measuring the Magic of Mutual Gaze with performance artist Marina Abramovic: she has used eye gaze as a tool for human connectedness throughout her work, such as The Artist is Present.

#### +/- DO PERSONALITY AND MENTAL STATE PREDICT YOUR BRAINWAVE SYNCHRONY?

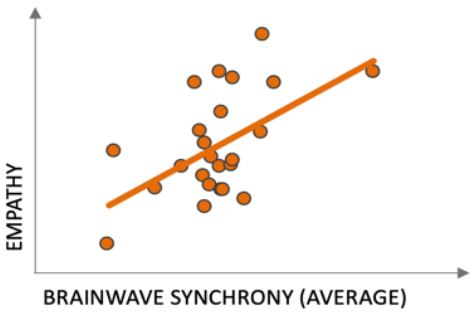


As can be seen in the scatterplot, pairs who were more empathetic on average, tended to also show higher brain synchrony. In this plot, each dot shows the average brainwave synchrony for all pairs that were similarly empathetic on average.

This has been a pretty consistent finding in our work. For example, more empathetic high school students exhibit more synchrony with their peers during class.

We also saw that synchrony increased for those pairs who remained more focused. This could mean that shared attention is a requirement for brain synchrony: for example, in other research we found that the more high schoolers attend to class together, and the more they attend to each other, the more their brainwaves synchronize.

<embedded video>

#### +/- DOES INTERACTING WITH EACH OTHER AFFECT YOUR MOOD AND SOCIAL BOND?

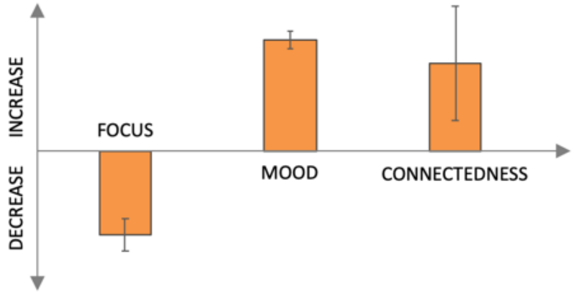


The bar graph shows how people rated their focus, mood and social connectedness to the other person before vs. after participating (i.e., for each self-report measure, you can see if it either increased or decreased).

First, people felt less focused after than before the experience. This could mean that people start out super focused and motivated, but that this ebbs over time. This is also common in regular laboratory experiments, where participants sometimes even fall asleep during the task (especially when they're lying down). Perhaps more meaningful is that people's mood improved and that most people felt closer to each other after participating than at the start of the experience. Unsurprisingly, this increase in self-reported social closeness was most prominent for people who didn't know each other well. These findings relate to previous work showing that face-to-face social interaction makes people happy and healthy, and tightens social bonds.

**+/- DOES THE "NEUROFEEDBACK" ENVIRONMENT AFFECT SYNCHRONY?**

We wanted to know if showing participants their EEG correlations in real time ("neurofeedback") might have a positive effect on their experience: Do you feel more connected to the other person? Does synchrony-based neurofeedback *increase* your brain synchrony? We also asked whether the feedback itself was 'meaningful': does it work only if it is real? Or is what matters whether people *believe* that what they see tells them something about how well they are connecting?

As it turns out, seeing is believing: Brain synchrony increased over time if participants were told that the audiovisual environment reflected their ongoing brain synchrony, even if the feedback was 'fake'. If you are interested in the notion of neurofeedback, see some discussions here and here.

**CREDITS & REFERENCES**

The Mutual Wave Machine is a collaborative project between the general public, the art/science collective DIKKER + OOSTRIK (Matthias Oostrik and Suzanne Dikker), researchers (most notably Georgios Michalareas, who led the offline synchrony analyses), artists (e.g., Peter Burr), our producers Danielle Boelling and Dana Bevilacqua, and several organizations (e.g., NWO and the Max Planck Institute for Empirical Aesthetics). Photographs included here were taken by Talia Herman, Sandra Kaas, and Lexus Hybrid Art.

<expandable sections collapsed>

**+ FULL CREDITS**

**+ RELATED ARTWORKS**

**+ RELATED RESEARCH**

<expandable sections expanded>

**+/- FULL CREDITS**

Matthias Oostrik, Suzanne Dikker, Georgios Michalareas, Peter Burr, Danielle Boelling, Melda Kahraman, Dana Bevilacqua, Marina Abramovic, Amalia Serafimaki, Matthew

Patterson-Curry, Lauren Silbert, Pandelis Diamantides, Marijn Struiksma, Phoebe Chen, Laura Noejovich, David Medine, Bogomir Doringer, Olof van Winden, Laura Gwilliams, Jean Jacques Warmerdam, David Poeppel, Annita Apostolaki, Shaista Dhanesar , Imke Kruitwagen, Eletta Daemen, Orsa Rebouskou, Stella Papazisi, Aspa Papazisi, Karlijn Blommers, Sascha Couvee, Ella Bosch, Jorik Geutjes, Irene Navarro, Alba Ortiz, Marta Regidor, Sandra Marquez, Belén Carrillo, Luna Miranda, Ester Abad, África Martín, Estefanía Granados, Iratxe de la Huerta, Isabel Rojo, Keumbee Lee, Patricia Plaza, Noelia Gómez, Brais Alen, and all the local volunteers

Netherlands Organisation for Scientific Research, VENI grant #275-89-018, EMOTIV, Universiteit Utrecht, NYU Poeppel Lab, Max Planck Institute for Empirical Aesthetics, Max Planck Center for Language, Music, and Emotion (NYU CLaME), TodaysArt, Lexus Hybrid Art, Marina Abramovic Institute, Pioneer Works, 3LD Art & Technology Center, NEON, Fundación Telefónica, Lowlands Festival (Lowlands Science), Nemo Science Center, EYE Filmmuseum, Benaki Art Museum, Creative Industries Fund NL, Nationale Wetenschapsagenda, De Hersenstichting, Stichting Niemeijer Fonds, FORMS Festival, Muse, Silicon Valley Contemporary

**+/-RELATED ARTWORKS**

- Measuring the Magic of Mutual Gaze
- The Compatibility Racer http://compatibilityracer.blogspot.com/
- Harmonic Dissonance: Synchron(icit)y
- Harmonic Dissonance
- The Artist is Present by Marina Abramovic
- Hive Mind by Produce Consume Robot & LoVid

**+/-RELATED RESEARCH**

- Brainwaves sync up in the classroom: original article here and here | selected press coverage here, here, and here
- Friends show the same brain activity: original article here | selected press coverage here
- Brainwave synchrony during pain perception: original article here | layman's blog here and here

#### MORE INFO

<expandable sections collapsed>

**+ BRAINWAVES: A SUPER SUPER BRIEF INTRO**

**+ A SUPER SUPER BRIEF INTRO TO EEG**

**+ TOOLS: QUESTIONNAIRES, EQUIPMENT, AND SOFTWARE**

**+ SCIENTIFIC ANALYSIS**

<expandable sections expanded>

#### +/- BRAINWAVES: A SUPER SUPER BRIEF INTRO

Our brain contains billions of neurons (latest estimate: 86 billion). A primary task of these neurons is to communicate information to each other and to the rest of our body via chemical signals. This process is accompanied by tiny electrical pulses. When lots of neurons fire together (like, when a musical tone reaches auditory cortex), this can be measured on the scalp as "brainwaves".

**Brainwaves** oscillate at different frequencies. Some of these brain rhythms have been specifically associated with different types of mental states: For example,

- **delta waves** (1-4 Hz, or: i.e., waves with an oscillation cycle of 1-4 times per second) are dominant during deep sleep.
- **theta waves** (~4-8 Hz) have been associated with focused attention and with language processing
- **alpha waves** (~8-12 Hz) are famous for their link to attentiveness and focus. They are quite easy to measure (for example when you close your eyes) and are often targeted in relaxation programs. It has also been suggested that people with AD(H)D have trouble 'controlling' their alpha waves.
- **beta waves** (~12.5 - 25 Hz) are very active during movement, like when you tap your fingers but also when you are merely thinking about tapping your fingers. Some researchers have argued that beta waves encode how concepts and objects are represented in our brains.
- **gamma waves** (~ 25 Hz and higher) are associated with many cognitive activities. Specifically relevant for this research is that gamma waves have been related to eye contact.

More about brainwaves

#### +/- A SUPER SUPER BRIEF INTRO TO EEG

**EEG (short for electroencephalography) is a tool used by researchers and doctors to measure brainwaves.** Sort of like a set of microphones, electrodes are placed on the scalp to listen in on the 'conversations' that take place inside our heads.
In the Mutual Wave Machine, we used the EMOTIV EPOC, which has 14 electrodes.
We are currently testing whether interpretable data during social interactions can also be acquired with the Muse, with 4 electrodes.

A big challenge in EEG research are so-called **motion artifacts**: the electrodes on the scalp don't just measure brain activity, they pick up *any* electric signal in our around the head. And muscular activity is quite electric… so much so that laughing, talking, even moving your eyes or blinking, will occlude brain activity. Some of this can be taken care of by filtering the signal, but not everything. To use the microphone analogy: say you're using microphones to record a conversation. They will pick up any sound in the room. Some sounds or noise may not be in the same frequency as the people's voices, so you can filter that out of the signal to hear better what people were saying. But if someone is screaming through a conversation, it is really difficult to ever recover what was said.

Also just like in the case of microphones, from an EEG recording it is much easier to **reconstruct *when* a brain response happened** than *where* it came from. In other words, we won't be able to conclude anything from the Mutual Wave Machine about which brain regions were involved, but we *can* say something about whether people's brain rhythms were showing similar patterns. And that's exactly what we did.

More about EEG

#### +/- TOOLS: QUESTIONNAIRES, EQUIPMENT, AND SOFTWARE

We used questionnaires that have been very well-validated. For example, the Inclusion of the Other in the Self Scale measures how close people feel to each other, the PANAS-X measures your mood, and the Interpersonal Reactivity Index measures how you react in different social situations. We then asked how each of these responses related to pairs' brainwave synchrony.

To collect the EEG data, we used the EMOTIV EPOC, an EEG headset with 14 electrodes. We built software that allowed us to record data from multiple headsets onto a single computer, and we then used openframeworks to analyze and visualize brainwave synchrony in real-time.

We are currently testing whether interpretable data during social interactions can also be acquired with the Muse, a headset with 4 electrodes. We built an iOS app to collect data from the Muse and then transmit it to a mother computer using Lab Streaming Layer. We then use python to compute brainwave synchrony, which is visualized with openframeworks.  This is a collaborative effort with David Medine, Jean-Jacques Warmerdam, Laura Gwilliams, and Phoebe Chen. The protocol will be posted to the BCI plus site.

We worked with Peter Burr on the aesthetics of the visuals.

#### +/- SCIENTIFIC ANALYSIS

After data collection, Georgios Michalareas wrote a series of MATLAB scripts that clean and analyze data using the FieldTrip toolbox.

- First, we discarded datasets with too many motion artifacts or other recording problems.
- Then, we cleaned the remaining datasets: this meant removing stretches of data with a lot of noise and electrodes that didn't have a good enough connection to the scalp to record brainwaves properly.
- The data was then filtered into different brainwave frequencies
- Then, for each frequency, we computed a few different types of brainwave synchrony. These types largely fall into two different categories: (1) do pairs show similar fluctuations in their brain states? (2) do pairs' brainwaves show the same shape?
- Finally, brainwave synchrony was compared between groups (for example the feedback vs. no feedback groups) and as a function of participants' responses to the questionnaire questions (for example, empathy).
